# *Lactiplantibacillus plantarum* PS128 restores microbiota-driven polyamine homeostasis to alleviate autism-like behaviors in *Fmr1* knockout mice

**DOI:** 10.64898/2025.12.15.694366

**Authors:** Yingmin Cui, Yingxuan Chen, Yen-Wenn Liu, Shan Zheng, Jialong Cui, Tianan Li, Ying-Chieh Tsai, Shanshan Gu, Ji Ma, Wei Zhang, Haipeng Li, Xiao-Bing Yuan, Yi-Hsuan Pan

**Author notes:** Correspondence (Y.H.P.), (X.B.Y.). These authors contributed equally (Yingmin Cui, Yingxuan Chen, Yen-Wenn Liu).

## Abstract

**Background:** Autism spectrum disorder (ASD) is biologically heterogeneous and has limited mechanism-informed interventions targeting core behavioral symptoms. Emerging evidence suggests that gut microbial metabolism shapes neurobehavioral outcomes, yet the specific metabolic pathways linking gut ecology to brain function remain incompletely understood. One candidate pathway is polyamine metabolism, which links microbial amino acid metabolism to host regulation and represents a plausible but underexplored contributor to ASD-related phenotypes. Notably, the psychobiotic *Lactiplantibacillus plantarum* PS128 has been reported to improve social behaviors in individuals with ASD, although the molecular basis for these effects is unclear. In this study, we used Fragile X mental retardation 1 knockout (*Fmr1* KO) mice, a well-established ASD model, to investigate microbiota-dependent metabolic mechanisms underlying autism-like behaviors.

**Results:** We found that autism-like behaviors in *Fmr1* KO mice were associated with gut dysbiosis, impaired intestinal barrier integrity, disruption of the arginine–ornithine–polyamine pathway, and elevated putrescine in the prefrontal cortex (PFC). Supplementation with PS128 remodeled the gut microbiota, reduced inflammation- and disease-associated taxa, improved intestinal structure and permeability, and restored polyamine homeostasis. Targeted metabolomics revealed an increased PFC-to-serum putrescine ratio in KO mice, which was normalized following PS128 intervention. This correction was accompanied by reduced expression of polyamine transporters in the PFC, including ATP13A family members. Causal experiments supported a functional role for putrescine, as peripheral elevation of putrescine induced autism-like behaviors in wild-type mice, whereas pharmacological inhibition of putrescine synthesis ameliorated behavioral deficits in *Fmr1* KO mice.

**Conclusions:** These findings identify putrescine metabolic dysregulation as a key contributor to autism-like phenotypes and support the existence of an ASD subtype defined by disruption of the arginine–ornithine–polyamine axis. Integrated multi-omics and causal perturbation analyses support a model in which microbiota-targeted intervention rebalances systemic polyamine regulation along the gut–brain axis, thereby improving ASD-relevant behaviors. Our work provides mechanistic evidence linking microbial metabolism to neurochemical homeostasis and highlights the translational potential of metabolic stratification in ASD.

**Graphical Abstract:** We identify dysregulated microbiota-driven polyamine homeostasis as a causal contributor to autism-like behaviors in *Fmr1* knockout mice. A psychobiotic intervention restores gut–brain metabolic balance and rescues social and repetitive behavioral deficits through coordinated microbial and host transport regulation.

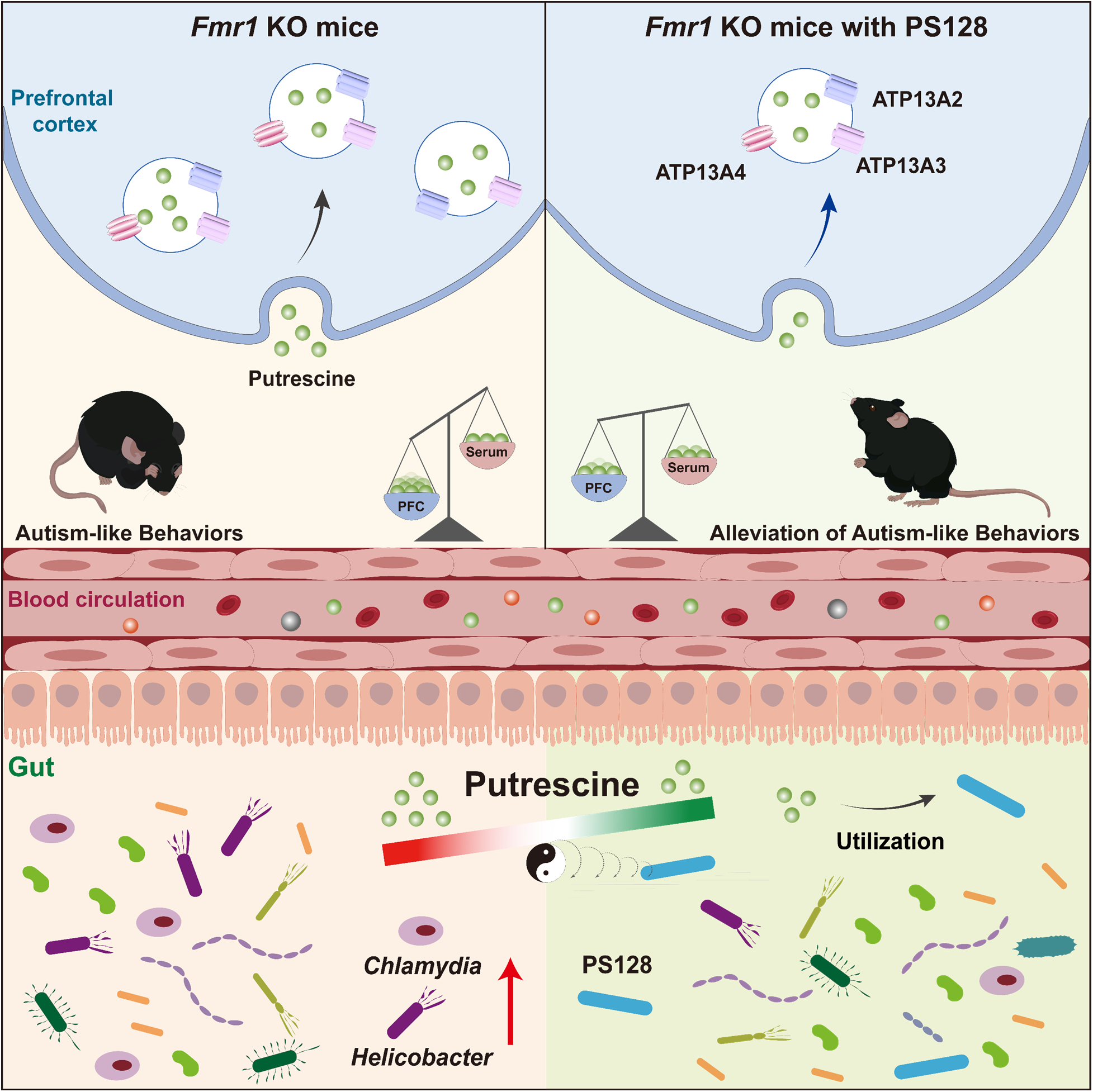

## Background

Autism spectrum disorder (ASD) encompasses a group of neurodevelopmental conditions characterized by deficits in social interaction and communication, accompanied by repetitive and restricted behaviors and interests. Both genetic and environmental factors contribute to ASD, with genetic influences playing a predominant role [1]. In recent years, ASD has emerged as a major public health concern due to its rising global prevalence [2, 3]. However, effective therapies targeting the core symptoms of ASD remain limited [4]. Growing evidence implicates metabolic disturbances in ASD pathophysiology, particularly involving amino acid metabolism, including branched-chain amino acids (BCAAs), arginine, and ornithine [5–8].

Microbial metabolites are increasingly recognized as key modulators of host physiology, with several shown to influence brain function through the gut–brain axis [9, 10]. Among the diverse classes of host- and microbiota-derived metabolites, putrescine is a polyamine primarily derived from arginine and ornithine catabolism. Putrescine interacts electrostatically with nucleic acids, proteins, lipids, and proteoglycans, thereby regulating gene expression, oxidative stress, cell proliferation, and inflammation [11]. Polyamine homeostasis is tightly regulated, and the intestinal microbiota represents a major exogenous source [12]. Notably, elevated levels of putrescine and its derivative spermine have been detected in the saliva of children with ASD [13], supporting a potential contribution of polyamine imbalance to ASD pathophysiology.

Probiotics, particularly psychobiotics, have attracted increasing interest as safe and accessible interventions capable of reshaping the gut microbiome and modulating neurobehavioral outcomes [14]. Clinical evidence indicates that daily supplementation with *Lactiplantibacillus plantarum* PS128 (PS128) has been associated with improvements in social behaviors and attention-related symptoms in individuals with ASD, with minimal adverse effects [15, 16]. However, the molecular pathways through which PS128 acts along the gut–brain axis to ameliorate ASD symptoms remain poorly defined. Fragile X mental retardation protein (FMRP), an RNA-binding protein encoded by the *Fmr1* gene, represents the most common monogenic cause of ASD [17], and its deficiency leads to both gastrointestinal and behavioral abnormalities in humans and mice [18, 19]. Accordingly, the *Fmr1* knockout (KO) mouse [20] provides a well-established model for dissecting gut–brain interactions in ASD and for uncovering the molecular mechanisms by which microbiota-targeted interventions may exert biological effects.

In this study, we investigated whether microbiota-targeted intervention can ameliorate autism-like phenotypes in *Fmr1* knockout mice. We identified disruption of the arginine–ornithine–polyamine metabolic pathway in the gut of KO mice and found that intervention with *Lactiplantibacillus plantarum* PS128 reshaped the gut microbiota and rebalanced peripheral–central putrescine homeostasis. This effect was accompanied by enrichment of beneficial bacterial taxa and suppression of inflammation- and disease-associated taxa in the gut, together with reduced expression of putrescine transporters in the prefrontal cortex (PFC). Furthermore, bidirectional manipulation of putrescine levels demonstrated a causal contribution of putrescine to autism-like phenotypes. These findings suggest putrescine as a candidate metabolic biomarker and therapeutic target for ASD.

## Results

### Microbiota-targeted intervention alleviates autism-like behaviors in *Fmr1* KO mice

To determine whether microbiota modulation mitigates behavioral abnormalities in *Fmr1* knockout (KO) mice, KO mice and their wild-type (WT) littermates received PS128-containing jelly every other day through voluntary oral consumption (Video S1). Behavioral performance was assessed using a comprehensive battery, including the open-field test (OFT), elevated plus maze (EPM), O-maze test (OMT), novel object recognition test (NORT), reciprocal social interaction test (RSIT), balance beam test (BBT), olfactory habituation/dishabituation test (OHDT), and tube test (TT) throughout the treatment period. These assays were selected to capture ASD-relevant behavioral domains, including anxiety-related behavior, repetitive and stereotypic behaviors, social interaction and novelty preference, and motor coordination.

Across most behavioral assays, PS128 intervention had minimal effects on WT mice (Figs. 1 and S1), and the behavioral alterations were primarily observed in *Fmr1* KO mice. In the OFT (Figs. 1A and S1A), KO mice exhibited significantly reduced total and central exploration distances compared with WT controls (Fig. 1B and 1C), indicating increased anxiety-like behavior. Following intervention, these measures were restored toward WT levels (Fig. 1A–1C), and time spent in the center zone increased (Fig. S1B and S1C), indicating attenuation of anxiety-like behavior in KO mice. No significant differences were detected in the EPM or OMT (Fig. S1D–S1I), suggesting that anxiety-related abnormalities in KO mice are context dependent.

**Fig. 1.**
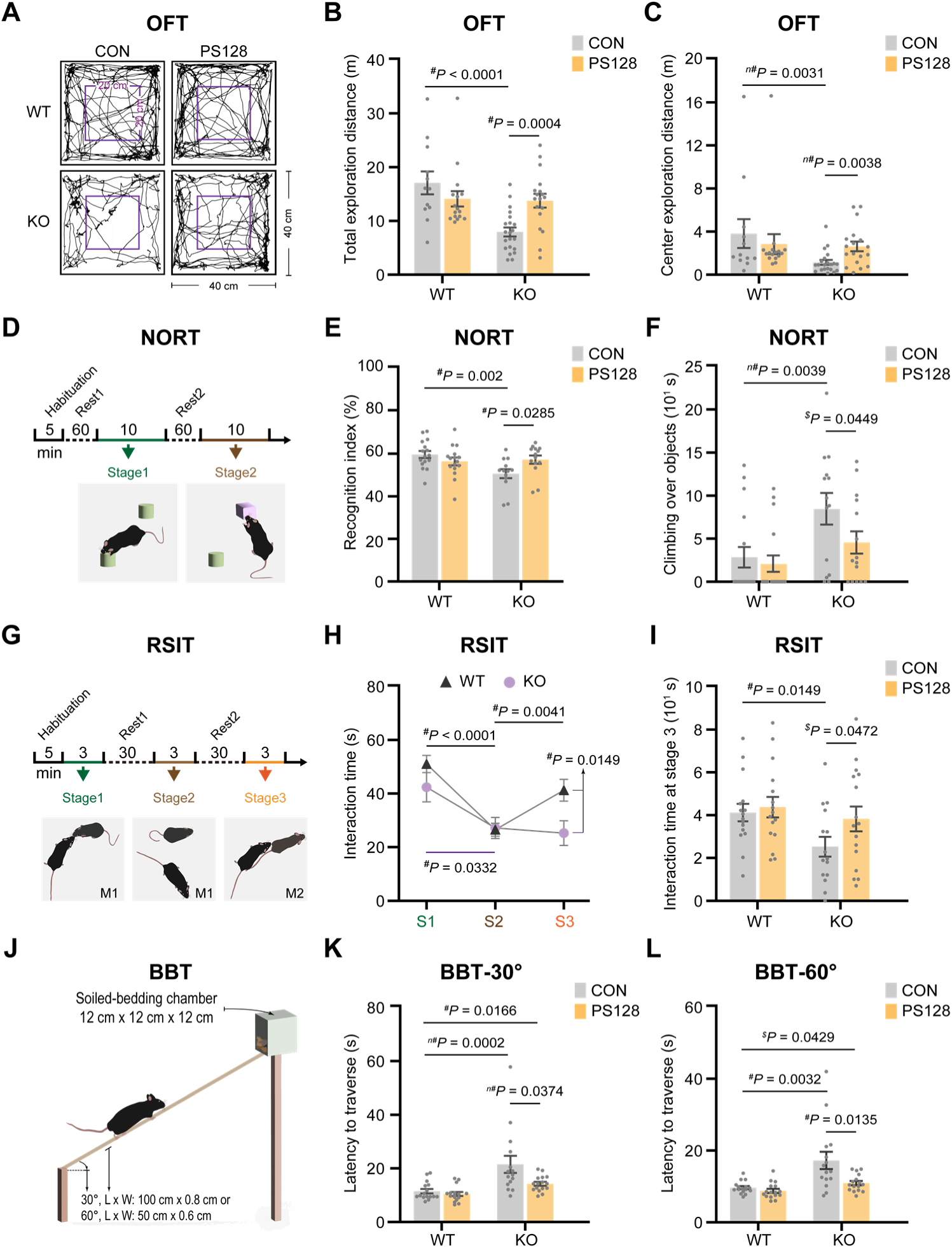
Microbiota-targeted intervention alleviates autism-like behaviors in *Fmr1* KO mice. **A–C** Representative traces in the open-field test (OFT) (**A**), total exploration distance (**B**), and center exploration distance (**C**), *n* = 12–21. **D–F** Schematic of the novel object recognition test (NORT) (**D**), recognition index (%) = (novel) / (novel + familiar) x 100, based on sniffing time (**E**), and total time spent climbing over objects (**F**), *n* = 13–18. **G–I** Schematic of the reciprocal social interaction test (RSIT) (**G**), interaction time with familiar or novel mice across three stages (**H**), and interaction time at Stage 3 (**I**), *n* = 15–16. S, stage. **J–L** Schematic of the balance beam test (BBT) (**J**), latency to traverse a 30° beam (**K**), or a 60° beam (**L**), *n* = 15–16. Data are presented as mean ± S.E.M. and gray dots indicate individual mice. One-tailed (*^$^*) or two-tailed (*^#^*) Student’s *t*-test and two-tailed (*^n#^*) Mann–Whitney U test were used where appropriate. A *P* value < 0.05 was considered statistically significant. See also Fig. S1.

In the NORT (Fig. 1D), KO mice showed a reduced recognition index (Fig. 1E) and excessive climbing on both familiar and novel objects (Fig. 1F), reflecting repetitive and stereotypic behaviors [21]. Similar repetitive climbing behavior was also observed in the OHDT (Fig. S1J–S1N, Videos S2 and S3), where KO mice frequently climbed and remained on the swab (Fig. S1J). Following intervention, both recognition deficits and stereotypic behaviors were ameliorated, with the recognition index restored toward WT levels and a marked reduction in excessive climbing.

In the RSIT (Fig. 1G), WT and WTPS mice displayed comparable interaction patterns (Fig. S1O). WT mice preferentially interacted with a novel mouse (Stage 3) over a familiar one (Stage 2), whereas KO mice did not (Fig. 1H and 1I), indicating impaired social novelty preference. Intervention significantly improved social novelty preference and recognition performance in KO mice (Fig. 1D–1I).

In the BBT (Fig. 1J), KO mice required more time to traverse both 30° and 60° beams (Fig. 1K and 1L), indicating impaired motor coordination. Traversal performance was improved following intervention in KO mice, although performance did not fully reach WT levels. In the TT (Fig. S1P), PS128-treated KO mice exhibited a higher probability of forward movement against unfamiliar KO or WT opponents (Fig. S1Q and S1R), suggesting enhanced social confidence. Collectively, microbiota-targeted intervention robustly improved core autism-like phenotypes in *Fmr1* KO mice (Figs. 1 and S1).

### Microbiota remodeling reduces inflammation- and disease-associated taxa in *Fmr1* KO mice

To characterize microbiota remodeling associated with behavioral improvement in *Fmr1* KO mice, we performed 16S rRNA sequencing and metagenomic profiling on fecal samples, together with fluorescence imaging of gut tissues, collected from KO mice and WT littermates before and during microbiota-intervention with PS128.

Before PS128 supplementation, 1-month-old KO mice (KO-1M) exhibited lower α-diversity than age-matched WT littermates (WT-1M) (Fig. 2A and 2B) and a distinct β-diversity profile (Fig. 2C). WT-1M mice harbored several beneficial genera, including *Bifidobacterium*, *Anaerostipes*, and *Blautia* (Fig. 2D), which are known to promote mucus secretion, protect against enteric pathogens, and modulate mucosal immunity [22]. In contrast, KO-1M mice displayed increased abundances of *Helicobacter* and *Chlamydia* (Fig. 2D), pathogenic taxa previously associated with gastrointestinal inflammation and neurobehavioral abnormalities [23, 24].

**Fig. 2.**
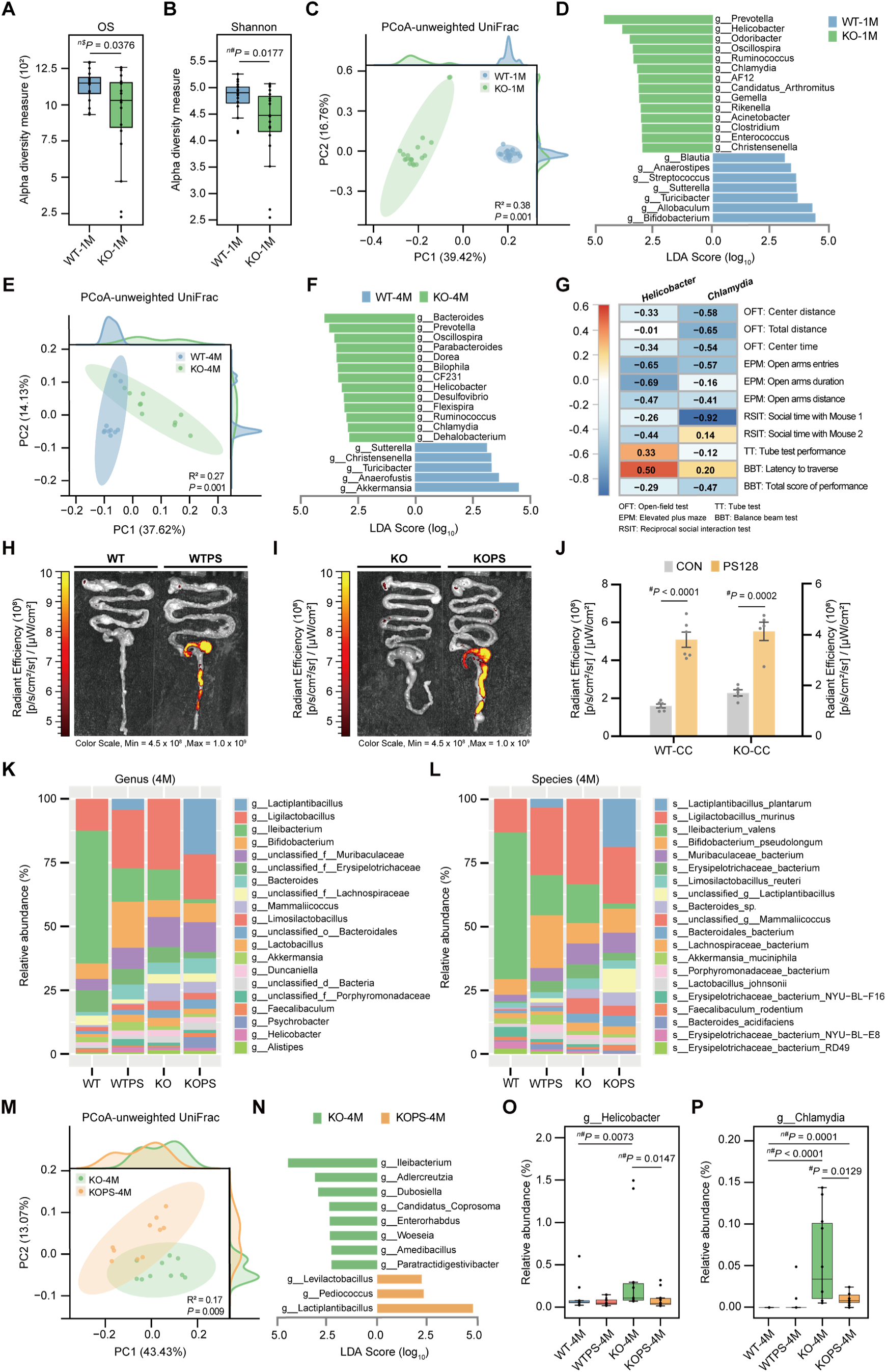
Microbiota-targeted intervention reshapes gut microbial community structure in *Fmr1* KO mice. **A–D** Gut microbiota analysis of 1-month-old *Fmr1* KO mice and their wild-type (WT) littermates without PS128 intervention. Observed species (**A**). Shannon diversity index (**B**). Principal coordinates analysis (PCoA) plot based on unweighted UniFrac distances (**C**). Histogram of linear discriminant analysis (LDA) scores from LEfSe analysis at the genus level, logarithmic LDA score cutoff > 3.0, *P* < 0.05 (**D**), *n* = 19–20. **E** and **F** Gut microbiota analysis of 4-month-old *Fmr1* KO mice and their WT littermates following 3 months of PS128 supplementation. PCoA plot (**E**). Histogram of logarithmic LDA scores, cutoff > 2.5, *P* < 0.05 (**F**), *n* = 10. **G** Spearman’s rank correlation analysis between the relative abundance of *Helicobacter* and *Chlamydia* and behavioral performance in 4-month-old *Fmr1* KO mice, *n* = 10–11.**H–J** Biodistribution of PS128 in the gastrointestinal tract. Representative ex vivo fluorescence images of the gut from *Fmr1* KO mice and their WT littermates after oral gavage with fluorescently labeled PS128 (**H** and **I**). Average radiant efficiency quantified from the cecum and colon (CC) (**J**), *n* = 5–6. **K** and **L** Relative abundance of gut microbiota at the genus (**K**) and species (**L**) levels in 4-month-old groups, *n* = 11–13. **M** and **N** Gut microbiota analysis comparing 4-month-old *Fmr1* KO mice with or without PS128 treatment (KO or KOPS). PCoA plot (**M**). Histogram of logarithmic LDA scores, cutoff > 2.2, *P* < 0.05 (**N**), *n* = 10-11. **O** and **P** Relative abundance of *Helicobacter* (**O**) and *Chlamydia* (**P**) across 4-month-old groups, *n* = 10. Data are presented as mean ± S.E.M. and each dot represents an individual mouse. Statistical significance was determined by two-tailed Student’s *t*-test (*^#^*), one-tailed (*^n$^*), or two-tailed (*^n#^*) Mann–Whitney U test, as appropriate. A *P* value < 0.05 was considered significant. See also Figs. S2 and S3.

By four months of age, KO mice (KO-4M) maintained reduced α-diversity (Fig. S2A and S2B) and distinct β-diversity relative to WT-4M mice (Fig. 2E), along with persistently elevated *Helicobacter* and *Chlamydia* (Fig. 2F). Spearman correlation analysis further revealed that higher abundances of these taxa were associated with more severe behavioral deficits in KO-4M mice (Fig. 2G), including reduced social interaction time (RSIT), prolonged traversal latency (BBT), and impaired tube test (TT) performance.

To enable in vivo tracking, CFDA/SE-labeled PS128 was first validated for growth and tolerance to simulated gastric and intestinal fluids and showed no detectable differences from unlabeled bacteria (Fig. S3A–S3C). Fluorescence imaging revealed preferential retention and colonization of the administered strain (PS128) in the cecum and colon of both WT and KO mice (Figs. 2H–2J and S3), as well as in the stomach and small intestine of WT mice (Fig. S3D, S3F, and S3N). Independently, analysis of fecal samples from mice receiving unlabeled PS128 supplementation demonstrated pronounced microbiota remodeling (Figs. 2K, 2L, and S2C–S2E). Following intervention, KO mice (KOPS-4M) exhibited α-diversity comparable to WT-4M littermates (Fig. S2F and S2G) and a distinct β-diversity profile relative to untreated KO-4M mice (Fig. 2M). At the taxonomic level, KOPS-4M mice showed enrichment of *Lactiplantibacillus* and *Bifidobacterium*, together with reductions in *Ileibacterium*, *Enterorhabdus*, *Helicobacter*, and *Chlamydia* (Figs. 2K, 2L, 2N–2P and S2H–S2M). Overall, these results demonstrate that microbiota-targeted intervention with PS128 robustly reshapes gut microbial community structure in *Fmr1* KO mice by reducing inflammation- and disease-associated taxa while enriching commensal genera, including *Lactiplantibacillus* and *Bifidobacterium*, in parallel with behavioral improvement (Figs. 2 and S2).

### Restoration of gut barrier integrity accompanies microbiota remodeling

To determine whether microbiota remodeling was associated with restoration of intestinal structure and function, we performed anatomical, histological, and physiological analyses, using WT littermates as references.

KO mice displayed significantly elevated intestinal permeability in the intestinal permeability assay (IPA), which was attenuated following microbiota-targeted intervention with PS128 (Fig. 3A and 3B). In addition, KO mice exhibited pronounced gastrointestinal abnormalities, including elongation of the cecum and colon (Fig. 3C–3E), distorted crypt architecture (Fig. 3F and 3G), and reduced goblet cell numbers (Fig. 3H–3J), as well as diminished MUC2-positive areas (Fig. 3K–3M). This intervention alleviated these abnormalities, partially restoring mucosal architecture and improving epithelial barrier integrity. Collectively, these results demonstrate that microbiota-targeted intervention with PS128 substantially improves intestinal barrier structure and function in *Fmr1* KO mice.

**Fig. 3.**
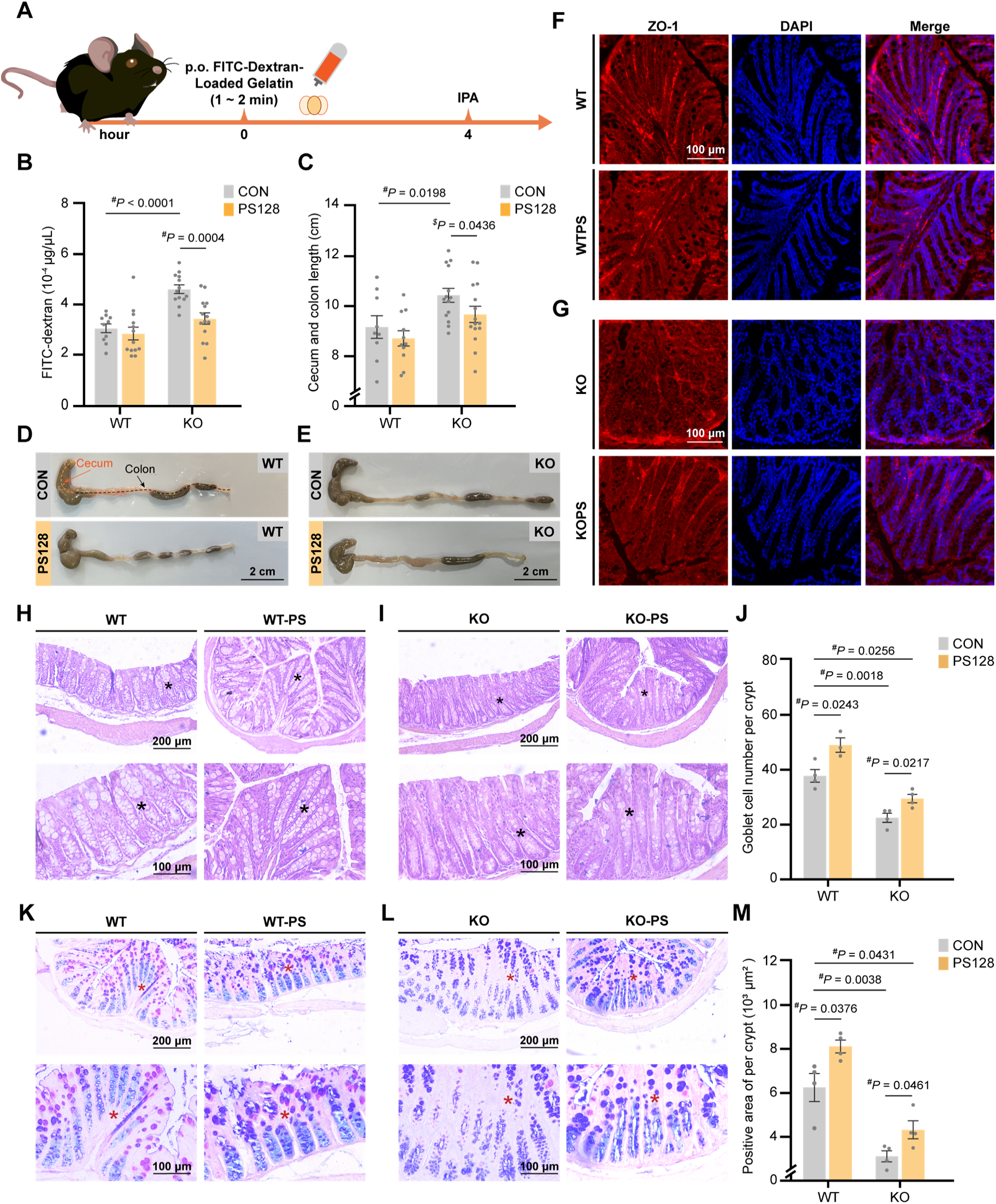
Microbiota remodeling restores intestinal barrier integrity in *Fmr1* KO mice. **A** and **B** Assessment of intestinal permeability. Schematic of intestinal permeability assay (IPA) (**A**). Serum FITC-dextran concentrations after orally self-consumed FITC–dextran–loaded gelatin (**B**), *n* = 10–14. **C–E** Gross anatomical assessment of the large intestine. Average length of the cecum and colon, *n* = 9–15 (**C**). Representative images of dissected tissues (**D** and **E**). **F** and **G** Colon integrity. Representative immunofluorescence images of colonic sections stained for ZO-1 (red) and nuclei (DAPI, blue). Scale bar, 100 µm. **H–J** Histological evaluation of colonic architecture. Representative H&E–stained sections of *Fmr1* KO and their WT littermates with/without PS128 intervention (**H** and **I**). Quantification of goblet cells per crypt (average of 3 crypts per section, 3 sections per mouse), *n* = 3–4 (**J**). Scale bars, 200 µm and 100 µm. **K–M** Analysis of mucin production. Representative AB–PAS–stained sections showing acidic and neutral mucins in blue and magenta for KO and their WT littermates with/without PS128 intervention (**K** and **L**). Quantification of MUC2-positive light blue and dark purple areas per crypt, 3 crypts per mouse, *n* = 4 (**M**). Scale bars, 200 µm and 100 µm. Data are mean ± S.E.M. and each dot represents an individual mouse. Statistical significance was determined by one-tailed (*^$^*) or two-tailed (*^#^*) Student’s *t*-test. A *P* value < 0.05 was considered significant.

### Modulation of the gut arginine–ornithine–polyamine pathway by microbiota-targeted intervention with *Lactiplantibacillus plantarum* PS128

To investigate the functional consequences of microbiota remodeling on gut metabolic pathways, we reanalyzed the metagenomic datasets and performed whole-genome sequencing of the administered strain (PS128) to assess its microbial metabolic potential in *Fmr1* KO mice.

Based on KEGG annotation, microbial functions in KO mice were primarily enriched in carbohydrate and amino acid metabolism (Fig. S4A). Among these categories, amino acid metabolism showed the most pronounced difference between KO and WT mice, with the highest LDA score (log_10_ = 3.55) (Fig. S4B). Microbiota-targeted intervention with PS128 further enhanced amino acid–related pathways in KO mice (Fig. S4C). These results highlight amino acid metabolic dysregulation as a prominent microbial feature associated with *Fmr1* deficiency and likely linked to physiological abnormalities.

We next reconstructed metagenome-assembled genomes (MAGs) by binning contigs derived from the metagenomic datasets [25] and identified gut–brain modules (GBMs) encoded by species-level genome bins (SGBs) (Fig. 4A). GBMs were categorized into eight functional clusters according to their relevance to gut–brain metabolic pathways. Among these, “Neurotransmitter metabolism I” (glutamate metabolism), “Amino acid metabolism II” (serine metabolism), and “Amino acid metabolism I” (arginine metabolism) contained the largest numbers of modules. Notably, “Amino acid metabolism I” showed the clearest difference between WT and KO mice and exhibited the most substantial recovery following microbiota-targeted intervention (Fig. 4B).

**Fig. 4.**
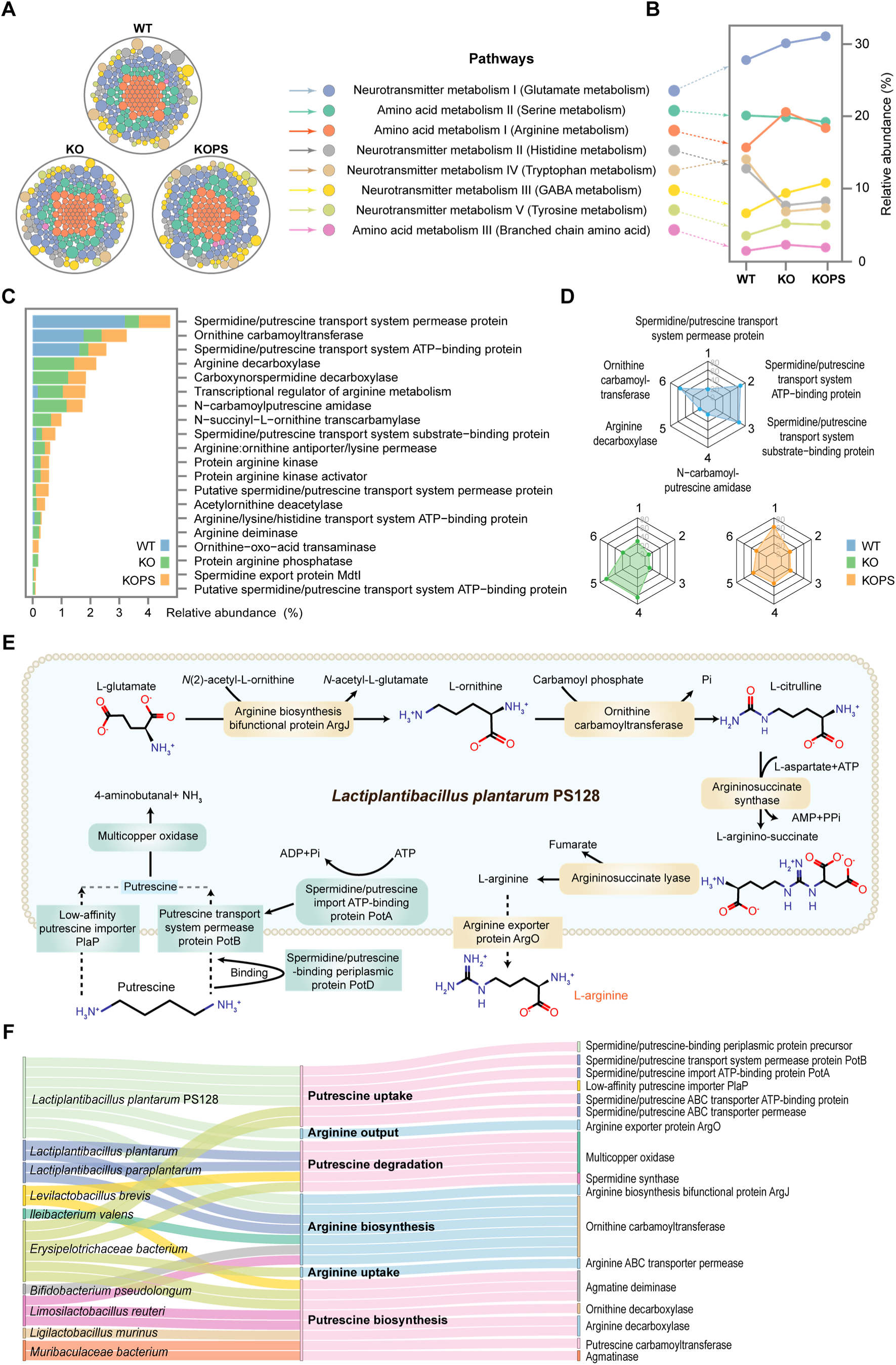
Microbiota-targeted intervention reprograms the gut arginine–ornithine–polyamine metabolic pathway. **A–D** Distribution of gut–brain modules (GBMs) at the species level. Each GBM is represented by a distinct color (**A**). GBMs were classified into eight functional clusters based on their association with gut–brain metabolic pathways. Relative abundance (%) of total modules within each GBM cluster (**B**). Relative abundance (%) of proteins encoded by bins related to “Amino acid metabolism I” (**C**). Key pathways in amine metabolism within “Amino acid metabolism I” (**D**). *n* = 11–13 per group. **E** Schematic of metabolic pathways annotated from whole-genome sequencing of PS128. **F** Sankey diagram showing correlations between genes encoding enzymes involved in arginine, ornithine, and spermidine/putrescine metabolism and bacterial strains enriched by PS128, together with the six most abundant bacterial species. See also Fig. S4.

Proteins encoded by bins within “Amino acid metabolism I” were annotated and quantified (Fig. 4C), and highly expressed proteins were visualized using radar plots (Fig. 4D). Compared with WT littermates, KO mice displayed reduced abundance of spermidine/putrescine transport ATP- binding and substrate-binding proteins, along with elevated levels of N-carbamoyl-putrescine amidase, indicating diminished bacterial uptake and increased microbial production of putrescine. KO mice also showed increased arginine decarboxylase and decreased ornithine carbamoyltransferase, reflecting upstream disruption of arginine and ornithine metabolism. Microbiota-targeted intervention with PS128 normalized these protein expression patterns, resulting in a more balanced amino acid and polyamine metabolic profile in KOPS mice (Fig. 4D).

Whole-genome sequencing of PS128 revealed genes involved in arginine biosynthesis and export, as well as in the uptake and degradation of spermidine and putrescine, whereas genes required for de novo polyamine biosynthesis were absent (Fig. 4E). This genomic profile is consistent with a putrescine-consuming role for PS128 in KO mice. In addition, functional annotation showed that PS128-enriched taxa, including *Lactiplantibacillus plantarum*, *Lactiplantibacillus paraplantarum*, and *Levilactobacillus brevis*, together with the six most abundant bacterial species, harbor complementary genetic capacities for arginine, ornithine, and polyamine uptake, degradation, and synthesis (Fig. 4F). Taken together, these analyses define a coordinated microbial network through which microbiota modulation following PS128 intervention shapes arginine–ornithine–polyamine metabolism in the gut of *Fmr1* KO mice.

### Peripheral–central putrescine imbalance is corrected by coordinated polyamine regulation

Behavioral improvement in *Fmr1* KO mice was most prominent in prefrontal cortex (PFC)–dependent social domains and striatum (STR)–associated repetitive behaviors (Figs. 1 and S1). Given that these behavioral changes coincided with remodeling of gut arginine–ornithine–polyamine metabolism (Figs. 4 and S4), we next examined whether peripheral metabolic alterations were reflected in polyamine homeostasis within the PFC and STR. To this end, we performed targeted metabolomic profiling of 40 neuroactive substances, seven short-chain fatty acids (SCFAs), and four additional fatty acids in both brain regions (Tables S1 and S2).

Volcano plot analysis revealed a marked elevation of putrescine in the PFC of KO mice relative to WT littermates (Fig. 5A), which was significantly reduced following intervention with PS128 (Fig. S5A). This effect was largely region specific, as putrescine levels in the STR showed only modest group differences (Fig. S5B and S5C). No significant differences were detected among groups for the 11 fatty acids measured in either brain region (Fig. S5D–S5G). Because putrescine circulates systemically (Fig. 5B), we next quantified serum putrescine using targeted UHPLC–MS/MS and calculated the PFC-to-serum ratio. KO mice exhibited a significantly elevated ratio compared with WT controls (Fig. 5C), whereas intervention with PS128 largely normalized this imbalance (Fig. 5D), indicating a coordinated adjustment of peripheral and central putrescine distribution in *Fmr1* KO mice.

**Fig. 5.**
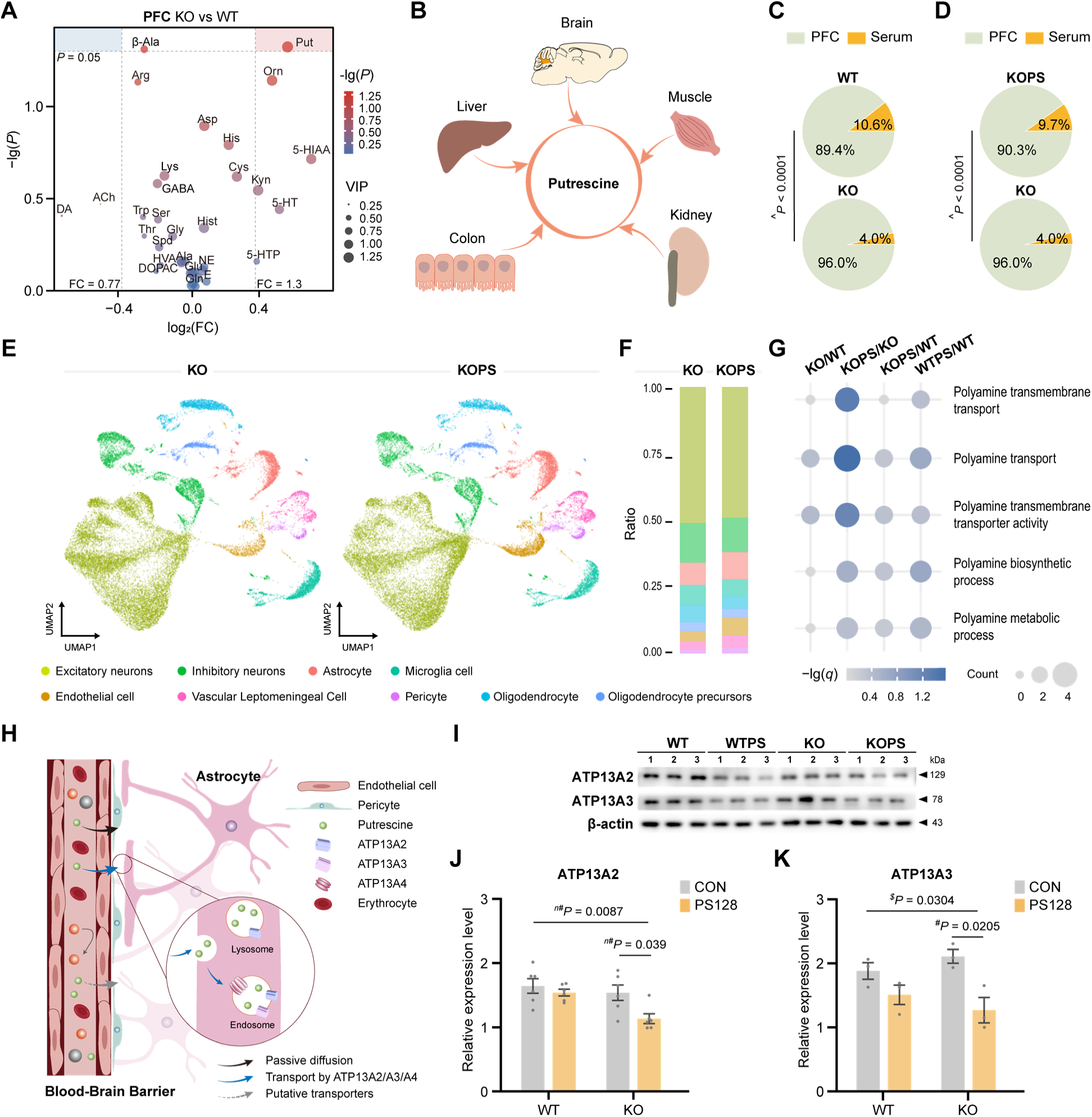
Microbiota-targeted intervention rebalances peripheral–central putrescine homeostasis and transporter expression. **A** Differential metabolites in the PFC between KO and WT mice. Colors indicate -log_10_(*P* value), and bubble sizes represent variable importance in projection (VIP) scores. The x-axis denotes log_2_(fold change). *n* = 3 biological replicates, each pooled from six individuals (18 mice per group). FC, fold change. **B** Schematic illustrating major peripheral organs as sources of circulating putrescine. **C** and **D** Ratios of putrescine in the PFC relative to serum in WT versus KO mice (**C**) and in KOPS versus KO mice (**D**). *n* = 6 mice per group. **E** and **F** UMAP projection (**E**) and stacked bar plots (**F**) of cell-type composition derived from single-nucleus RNA sequencing (snRNA-seq) of PFC from 15-week-old mice. *n* = 2 biological replicates per group, each pooled from 3–4 individuals (7–8 mice per group). **G** Gene Ontology (GO) enrichment of differentially expressed genes (DEGs) in the PFC, highlighting pathways related to polyamine transport. -log_10_(*q* value) >1.3 was considered significant. Count, number of DEGs. DEGs were defined as *P* < 0.05 and |log_2_ fold change| ≥ 0.12 with expression in at least 10% of PFC cells (Methods). **H** Schematic of two principal routes for putrescine transport. Putrescine crosses the blood–brain barrier (BBB) by passive diffusion or via active transport mediated by P5B-type ATPases (ATP13A2, ATP13A3, ATP13A4) and other putative transporters. **I–K** Western blot of ATP13A2, ATP13A3, and β-actin in the PFC (**I**), with densitometric quantification of ATP13A2 (**J**) and ATP13A3 (**K**) normalized to β-actin. *n* = 3–6 mice per group. Data are presented as mean ± S.E.M. and gray dot represents individual mice. Statistical significance was determined using one-tailed (*^$^*) or two-tailed (*^#^*) Student’s *t*-test, or two-tailed (*^n#^*) Mann–Whitney U test as appropriate. A chi-square (^) test was used in (**C** and **D**). A *P* value < 0.05 was considered significant. See also Figs. S5 and S6.

To explore molecular changes accompanying this adjustment (Fig. 5D), we performed single-nucleus RNA sequencing (snRNA-seq) on PFC tissue from KO and WT mice with or without PS128 treatment. Overall cell-type composition was not appreciably altered (Figs. 5E, 5F, and S6A). However, transcriptomic analysis revealed a predominance of downregulated genes following PS128 treatment (Fig. S6B). Downregulated genes were enriched for pathways related to chromosome organization, glycoprotein metabolism, and macroautophagy (Fig. S6C), whereas upregulated genes were associated with telomere organization and protein folding (Fig. S6D). Notably, pathways involved in polyamine transmembrane transport were selectively downregulated in PS128-treated KO mice (Fig. 5H).

These pathways included *Atp13a2*, *Atp13a3*, and *Atp13a4* (Fig. S6B and S6E), which encode mammalian P5B-type ATPases implicated in cellular polyamine handling. Reduced expression of ATP13A2 and ATP13A3 in PS128-treated KO mice (KOPS) was confirmed by Western blotting (Fig. 5I–5K). In addition, *Atp13a4*, which is enriched in astrocytes forming part of the blood–brain barrier (Fig. S6F), also exhibited reduced expression (Fig. S6G). These results indicate that microbiota-targeted intervention with PS128 downregulates polyamine transport–related pathways in the PFC while restoring peripheral–central putrescine equilibrium in *Fmr1* KO mice.

### Bidirectional manipulation of putrescine metabolism modulates autism-like behaviors in mice

To determine whether perturbation of putrescine metabolism contributes to autism-like behaviors, we administered putrescine dihydrochloride intraperitoneally (200 mg/kg) to 2-month-old wild-type (WT) mice (Fig. 6A). Although this treatment did not raise putrescine levels in the prefrontal cortex (PFC) (Fig. S7A), it was sufficient to induce multiple autism-like behavioral abnormalities. Treated WT mice showed reduced total exploration distance, center exploration distance, and center exploration time in the open-field test (OFT) (Fig. 6B–6F), impaired social interaction in the reciprocal social interaction test (RSIT) (Fig. 6G), and prolonged traversal time on the 30° balance beam (Fig. 6H). These results indicate that elevated peripheral putrescine (Fig. S7B) is sufficient to induce autism-like behavioral phenotypes in WT mice.

**Fig. 6.**
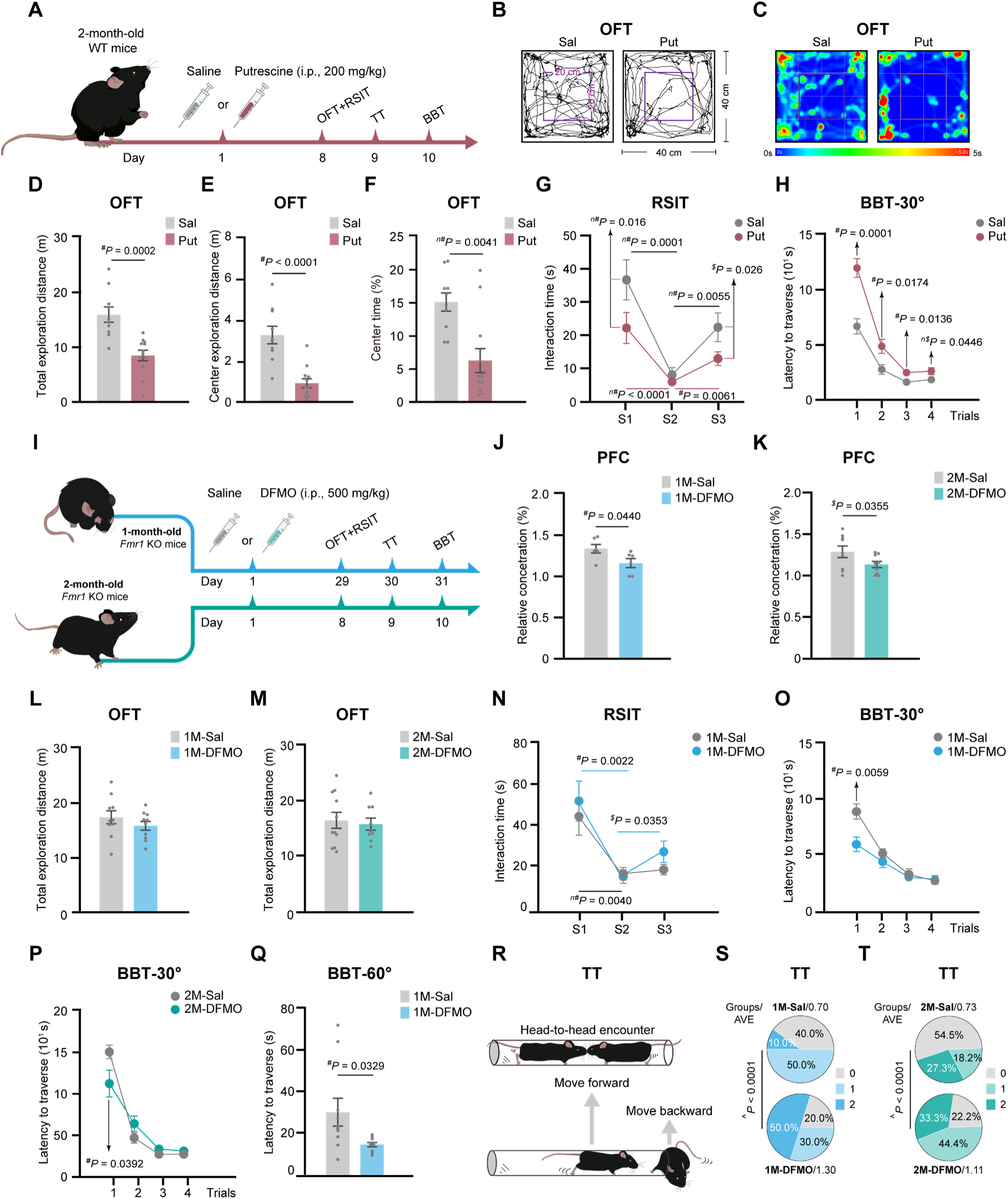
Putrescine imbalance causally modulates autism-like behaviors in mice. **A** Schematic of putrescine intervention in 2-month-old WT mice and behavioral tests. i.p., intraperitoneal injection. **B–F** Representative traces (**B**) and heatmap (**C**) of the open-field test (OFT), total exploration distance (**D**), center exploration distance (**E**), and time spent in the center area (**F**). **G** Interaction time with familiar or novel mice across three stages of the reciprocal social interaction test (RSIT). S, stage. **H** Latency to traverse a 30° beam across four trials of the balance beam test (BBT). Effects of exogenous putrescine intervention in 2-month-old WT mice (**B–H**), *n* = 10–12. **I** Schematic of DFMO intervention in 1- or 2-month-old *Fmr1* KO mice. i.p., intraperitoneal injection. **J** and **K** Relative putrescine concentration in the PFC after DFMO or saline injection from 1 month (**J**) or 2 months (**K**) of age, *n* = 6–9. **L** and **M** Total exploration distance in the OFT. **N** Interaction time with familiar or novel mice across three stages of the RSIT. **O–Q** Latency to traverse a 30° beam across four trials (**O** and **P**) or a 60° beam (**Q**) of the BBT. **R–T** Schematic of the tube test (TT) (**R**) and average score (AVE) per mouse obtained in the TT (**S** and **T**). Effects of endogenous putrescine inhibition by DFMO in 1- or 2-month-old *Fmr1* KO mice (**J–T**), *n* = 9–11. Data are presented as mean ± S.E.M. and gray dots indicate individual mice. One-tailed (*^$^*) or two-tailed (*^#^*) Student’s *t*-test, one-tailed (*^n$^*) or two-tailed (*^n#^*) Mann–Whitney U test were used as appropriate. A chi-square (^) test was used in (**S** and **T**). A *P* value < 0.05 was considered significant. See also Fig. S7.

We next examined whether reducing putrescine synthesis could ameliorate autism-like behaviors in *Fmr1* KO mice. To this end, we used α-difluoromethylornithine (DFMO), an irreversible inhibitor of ornithine decarboxylase (Fig. 6I). Intraperitoneal DFMO administration (500 mg/kg) significantly lowered PFC putrescine levels in both 1- and 2-month-old KO mice (Fig. 6J and 6K). DFMO had no significant effect on locomotor activity in the OFT (Fig. 6L and 6M); however, compared with 2-month-old KO mice, DFMO modestly reduced center exploration distance and time in 1-month-old KO mice (Fig. S7C–S7F), suggesting age-dependent effects on anxiety-like behavior. Despite this, DFMO treatment increased social interaction time in the RSIT (Figs. 6N and S7G) and shortened traversal time on the 30° beam during the first trial of the balance beam test (BBT) (Fig. 6O and 6P). Early DFMO intervention also improved performance on the 60° beam in KO mice (Fig. 6Q). Moreover, DFMO-treated KO mice exhibited tube test performance comparable to that of PS128-treated KO mice (KOPS) (Figs. 6R and S1R), achieving higher winning scores than untreated KO mice (Fig. 6S and 6T).

Altogether, these results demonstrate that elevated peripheral putrescine is sufficient to induce autism-like behaviors, whereas pharmacological suppression of putrescine production ameliorates these deficits, supporting a causal role for putrescine imbalance in behavioral abnormalities in *Fmr1* KO mice.

## Discussion

In this study, we show that altered microbial composition and dysregulated arginine–ornithine–polyamine metabolism are closely linked to autism-like behaviors in *Fmr1* KO mice. Microbiota-targeted intervention with *Lactiplantibacillus plantarum* PS128 reshaped the gut microbiota, improved intestinal integrity, and rebalanced putrescine homeostasis across the gut–brain axis, in part through coordinated regulation of microbial metabolism and host transport systems. Our data identify putrescine as a microbiota-responsive modulator of behavioral abnormalities and suggest that correction of the microbiota–polyamine–brain axis can occur through an integrated microbial–metabolic–neuronal pathway. These findings provide a conceptual framework for metabolic stratification and psychobiotic-based interventions in autism spectrum disorder.

Autism spectrum disorder (ASD) has substantial etiologic and phenotypic heterogeneity [4]. Genetic and environmental factors converge on the microbiota–gut–brain axis, where altered microbial metabolism influences amino acid turnover, neurotransmitter synthesis, epithelial integrity, and immune signaling [26]. Consistent with previous reports, *Fmr1* KO mice in our study exhibited repetitive and restricted behaviors, social impairments, and motor coordination deficits. These behavioral abnormalities were accompanied by intestinal dysbiosis, barrier dysfunction, and metabolic imbalance. Together, these features mirror reports of altered amino acid, lipid, and xenobiotic metabolism in ASD [27], as well as evidence linking mucosal barrier disruption [20] and microbial metabolite shifts to behavioral phenotypes [28, 29].

Sustained microbiota-targeted intervention with *Lactiplantibacillus plantarum* PS128 (PS128) markedly alleviated autism-like behaviors in *Fmr1* KO mice. Intervention with PS128 reduced the elevated abundances of *Helicobacter* and *Chlamydia*, two taxa highly correlated with autism-like behaviors in untreated *Fmr1* KO mice, and restored beneficial genera, such as *Lactiplantibacillus* and *Bifidobacterium*. While modulation of microbial composition is an anticipated outcome of probiotic intervention, we then asked whether such changes were accompanied by improvements in intestinal architecture or barrier-related features. Consistent with this possibility, intervention with PS128 ameliorated gastrointestinal abnormalities and barrier-associated features, supporting the conclusion that stabilization of microbial ecology contributes to improved intestinal integrity [30]. In addition, intervention increased the abundance of *Lactiplantibacillus paraplantarum*, *Levilactobacillus brevis*, and *Bifidobacterium pseudolongum*. Notably, *Lactiplantibacillus paraplantarum* has been reported to improve diabetes-related gastrointestinal complications and exert anti-inflammatory effects [31, 32]. These observations support a psychobiotic mode of action in which microbiota-targeted intervention promotes gut microbial homeostasis and intestinal integrity [15, 16, 33].

Beyond compositional rearrangement, intervention produced a distinct functional correction centered on the arginine–ornithine–polyamine metabolic pathway. Among polyamines, putrescine occupies a unique position at the interface of gut microbial metabolism and host regulation, rendering it sensitive to microbiota-driven metabolic perturbations [34]. Because the administered strain (PS128) lacks de novo putrescine biosynthesis but encodes uptake and catabolic functions, it likely contributes to the regulation of systemic putrescine handling rather than serving as a primary source of polyamine production. Consistent with this model, intervention with PS128 normalized the elevated prefrontal cortex (PFC)-to-serum putrescine ratio in *Fmr1* KO mice toward wild-type levels. This correction was accompanied by reduced expression of the polyamine transporters ATP13A2 and ATP13A3 in the PFC, together with decreased *Atp13a4* expression in astrocytes at the single-cell level in PS128-treated *Fmr1* KO mice. ATP13A2 and ATP13A3 are major determinants of mammalian polyamine uptake [35], and their downregulation is consistent with reduced prefrontal cortical accumulation of putrescine. Such coordinated changes suggest a homeostatic adjustment to altered polyamine availability, in line with their established roles in mammalian polyamine transport and cellular polyamine balance. Mutations or dysfunction in these transporters have been associated with ASD and related neurodevelopmental conditions [36–39]. These observations link microbial activity to central polyamine handling and support a model in which microbiota-targeted intervention modulates gut-derived putrescine flux and neuronal transporter expression.

We identify putrescine as a microbiota-sensitive regulator of autism-like behaviors. Elevating peripheral putrescine was sufficient to induce anxiety-like behaviors, social impairments, and motor deficits in wild-type mice, whereas inhibition of endogenous putrescine synthesis with DFMO ameliorated these abnormalities in *Fmr1* KO mice. At physiological pH, putrescine carries a positive charge and interacts with nucleic acids, phospholipids, and anionic proteins, thereby influencing macromolecular conformation, autophagy, membrane dynamics, and inflammatory signaling. Altered polyamine handling, including astrocytic uptake via ATP13A4, has been shown to modulate excitatory synaptic transmission and neuronal physiology [40]. Because putrescine serves as a metabolic precursor for astrocyte-derived γ-aminobutyric acid (GABA) through a noncanonical MAO-B- and ALDH-dependent pathway [41], perturbations in putrescine availability may reshape inhibitory tone and excitation–inhibition balance in cortical circuits. Clinical observations of elevated salivary putrescine [13] and increased activity of the gut microbial arginine–polyamine program in children with ASD [42] further support the concept that putrescine delineates a metabolically defined ASD subtype that may be preferentially responsive to microbiota-targeted interventions.

In our study, initiating DFMO at four weeks of age raised safety considerations, as juvenile *Fmr1* KO mice already exhibited mild anxiety-like behaviors. These results suggest that early inhibition of endogenous putrescine synthesis may exert age-dependent adverse effects on emotional regulation. Across behavioral endpoints, microbiota-targeted intervention produced broader and more consistent behavioral improvements than DFMO, consistent with its dual ecological (microbiota restructuring) and metabolic (polyamine modulation) actions.

Probiotic-based approaches are increasingly recognized as safe, non-pharmacological options for neurological and psychiatric disorders [43, 44]. *Lactobacillus reuteri* restored social behavior through vagus-dependent oxytocinergic and dopaminergic signaling [45]. The microbial metabolite 5-aminovaleric acid improved autism-like behaviors in BTBR mice by reducing pyramidal neuron excitability in the PFC [29]. *Bacteroides uniformis* reinstated excitation–inhibition balance in *Chd8^+/−^* mice by modulating intestinal amino acid transport and reducing prefrontal glutamine levels [46]. In a valproic acid–induced ASD mouse model, PS128 ameliorated social and memory deficits by enhancing oxytocin signaling and increasing dendritic complexity in the hippocampus and PFC [47]. Collectively, these studies support a framework in which psychobiotics influence brain function through coordinated effects on microbial metabolites, host transport systems, and cortical circuit activity. Because metabolic abnormalities have been widely reported in ASD across multiple metabolomics studies, future diagnostic strategies may complement current behavior-based assessments (e.g., DSM-5-TR) with metabolite-informed stratification.

This study has several limitations. We relied on a single genetic model, which limits the generalizability of our findings to other ASD subtypes. Although our multi-omics analyses support a gut-driven influence on polyamine homeostasis, the specific contributions of individual transporters and brain cell types remain to be delineated. Future studies incorporating transporter-specific perturbations, isotope-labeled tracing of gut-to-brain polyamine flux, and in vivo circuit-level measurements during behavior will help clarify mechanistic causality. Clinical translation will also require longitudinal studies integrating microbiome, metabolomic, and behavioral profiling in human cohorts. Ultimately, microbiota-targeted interventions may be combined with pharmacological or behavioral approaches and tailored to baseline metabolic phenotypes, enabling precision treatment strategies for ASD subtypes characterized by disrupted arginine–ornithine–polyamine metabolism.

## Conclusion

Our findings identify putrescine as a candidate metabolic marker for an ASD subtype and support a model in which *Lactiplantibacillus plantarum* PS128 modulates gut-derived polyamine metabolism and central polyamine transporter expression, thereby driving robust improvements in social and stereotypic behaviors.

## Methods

### Animals and Ethical Statement

Female C57BL/6J mice were obtained from Shanghai Jihui Laboratory Animal Care Co., Ltd. (Cat# SHJH-M-01002), and male *Fmr1* knockout (KO) mice (RRID: IMSR_JAX:004624) were obtained from The Jackson Laboratory. To generate *Fmr1* KO mice on a uniform C57BL/6J genetic background, the KO line was backcrossed with wild-type C57BL/6J mice for ten generations. Male wild-type and *Fmr1* KO littermates on a C57BL/6J background were used for all experiments. Mice were housed in groups of six under specific pathogen-free (SPF) conditions at the multifunctional platform for innovation (Platform 101). The animal facility was maintained at 22 ± 1°C and 50 ± 5% relative humidity under a twelve-hour light/twelve-hour dark cycle (lights on at 6:00 a.m. and off at 6:00 p.m.). All mice had ad libitum access to water and a standard chow diet, with daily food intake set at 2.5 g per mouse from 1 month of age and 3.0 g per mouse from 2 months of age onward.

### Bacterial Strains and Growth Conditions

*Lactiplantibacillus plantarum* PS128 was obtained from Centro Sperimentale del Latte Srl (Cat# G001723) and cultured in de Man-Rogosa-Sharpe (MRS) broth (Solarbio, Cat# M8540) at 37°C for liquid cultures or on MRS agar plates supplemented with 1.5% (w/v) agar (Sangon Biotech, Cat# A505255-0250).

### Preparation of Jelly Enriched with PS128, Blank Controls, and FITC

Gelatin-based jellies were prepared as follows. Briefly, 2.1 g of gelatin (Sinopharm, Cat# 10010326) and 0.3 g of sucrose (Sigma, Cat# V900116) were dissolved in 22.5 mL of sterilized double-distilled water (ddH_2_O) by heating to 55°C under constant stirring. The solution was then diluted with sterilized ddH_2_O prewarmed to 55°C to a final volume of 30 mL while continuously stirred.

The gelatin solution was evenly divided into two sterilized glass beakers and one 50-mL centrifuge tube (10 mL each) and maintained at 37°C. Lyophilized *Lactiplantibacillus plantarum* PS128 (0.46 g) was added to one beaker, and fluorescein isothiocyanate (FITC)–dextran (0.6 g, Aladdin, Cat# F121152) was added to the centrifuge tube. All procedures involving FITC were performed under light-protected conditions. Both beakers were placed on magnetic stirrers in a laminar-flow hood, whereas the centrifuge tube was mixed using a Genie 2 vortex mixer until fully homogenized. Aliquots of 150 µL from the PS128-enriched and blank gelatin solutions and 300 µL from the FITC–dextran solution were dispensed into sterilized custom-made 0.5-mL jelly molds. The molds (0.9 cm in diameter and 1.2 cm in height) were generated by trimming the upper portion of 2-mL cryogenic vials with conical round bottoms. The filled molds were incubated at 4°C for 2 hours to allow complete gelation.

After solidification, jellies were removed from the molds using sterilized tweezers. Each mouse was provided with one PS128-enriched jelly containing approximately 2.07 × 10^9^ colony-forming units (CFU) of *Lactiplantibacillus plantarum* PS128 once every two days. FITC-enriched jellies were prepared fresh as needed, and each mouse was provided with one jelly delivering 0.018 g of FITC–dextran per 30-g body weight.

### Training Mice for Voluntary Jelly Intake

Male wild-type (WT) and *Fmr1* knockout (KO) littermates at 3 weeks of age were housed separately for 1 week and then randomly assigned to four groups: WT, WTPS, KO, and KOPS. Mice in the WTPS and KOPS groups were provided with one PS128-enriched jelly every two days, whereas mice in the WT and KO groups were provided with one blank jelly on the same schedule (Video S1). To facilitate recognition and voluntary consumption of the jelly, the jelly surface was lightly coated with powdered standard chow during the first three feeding sessions.

### Behavioral Experiments

Behavioral testing was initiated after 28 consecutive days of exposure to PS128-enriched jelly or blank jelly. All tests were conducted between 9:00 a.m. and 3:00 p.m., with a minimum 2-day interval between consecutive tests to minimize stress. Prior to behavioral assessment, mice were handled for 10 minutes per day for 3 consecutive days and acclimated to the testing room for 2 hours on the test day.

Behavioral assays were performed in the following order: open-field test, novel object recognition test, elevated plus maze test, O-maze test, reciprocal social interaction test, olfactory habituation/dishabituation test, tube test, and balance beam test. Mouse behavior during training and testing sessions was recorded using the ANY-maze behavioral tracking system (version 7.20) and/or video cameras. All apparatuses were cleaned with 35% ethanol between individual mice, and a 5-minute interval was allowed between sessions to dissipate residual odors. Detailed procedures for each behavioral test are provided in Supplementary Material 1.

### Intestinal Permeability Assay

Four-month-old mice were fasted for 4 hours before actively consuming FITC–dextran–enriched jelly at a dose of 0.6 mg per g body weight. Four hours after jelly consumption, approximately 800 µL of venous blood was collected from each mouse via orbital blood sampling under isoflurane anesthesia and transferred to a 1.5-mL microcentrifuge tube. Blood samples were allowed to clot at 25°C for 1 hour, followed by centrifugation at 3,000 rpm for 10 minutes. The supernatant was carefully transferred to a clean 1.5-mL microcentrifuge tube and centrifuged again at 12,000 rpm for 10 minutes at 4°C.

A total of 200 µL of clarified serum was mixed with an equal volume of phosphate-buffered saline (PBS), and 150 µL of the diluted serum was added in duplicate to a 96-well flat-bottom black microplate. Fluorescence intensity of FITC–dextran in serum was measured using a spectrofluorometer at excitation and emission wavelengths of 485 nm and 528 nm, respectively. A standard curve was generated using serial dilutions of FITC–dextran in PBS (0, 1.0, 2.0, 4.0, and 6.0 × 10^-4^ µg/µL), and serum FITC–dextran concentrations were calculated by interpolation according to the manufacturer’s instructions.

### Histological Evaluation of Transverse Colon

Colon segments (1.5 cm in length, located 1.5–3.0 cm proximal to the distal colon) were excised and fixed in 4% paraformaldehyde in phosphate-buffered saline (PBS; pH 7.4) at 4°C for 24 hours. After fixation, tissues were washed in PBS, dehydrated through a graded ethanol series, and embedded in paraffin. Paraffin-embedded samples were sectioned at a thickness of 5 µm and mounted onto adhesive microscope slides.

Sections were deparaffinized, rehydrated, and stained with hematoxylin and eosin (H&E, Beyotime, Cat# C0105S) or Alcian blue–periodic acid–Schiff (AB–PAS, Leagene, Cat# DG0007). Images were acquired using an LED-illuminated light microscope. For quantitative analysis, three consecutive intact crypts per section, defined as fully visible and undamaged structures, were randomly selected. Measurements were performed on at least three independent sections per mouse.

### Immunofluorescence Assay

Paraffin-embedded sections were subjected to antigen retrieval by incubation in sodium citrate buffer (10 mM, pH 6.0) at 99°C for 15 minutes. Sections were then blocked in a solution containing 0.5% Triton X-100 (Sigma, Cat# T8787-100ML), 5% goat serum (Beyotime, Cat# C0265), and 5% bovine serum albumin (Sangon Biotech, Cat# A602440-0050) for 2 hours at room temperature. After blocking, sections were incubated with a primary antibody against ZO-1 (1:200, Thermo Fisher, Cat# ZO1-1A12) at 4°C for 36 hours. Following rinsing in phosphate-buffered saline (PBS), sections were incubated with an appropriate secondary antibody (1:500, Thermo Fisher, Cat# A-11003) for 2 hours at 37°C. Fluorescent signals were imaged using a confocal laser scanning microscope with excitation and emission wavelengths of 557 nm and 572 nm, respectively.

### Treatment Protocols for DFMO and Putrescine

α-Difluoromethylornithine (DFMO, MedChemExpress, Cat# HY-B0744) and putrescine dihydrochloride (Sangon Biotech, Cat# A600793) were dissolved in sterile saline at room temperature and administered via intraperitoneal injection (i.p.). For juvenile *Fmr1* knockout (KO) mice, DFMO treatment was initiated at 4 weeks of age at a dose of 500 mg per kg body weight, administered every other day for 28 days. After mice reached 2 months of age, DFMO administration was continued throughout the behavioral testing period until tissue collection. For adult mice, *Fmr1* KO mice received DFMO at a dose of 500 mg per kg body weight once daily for 7 consecutive days, whereas adult wild-type (WT) mice received putrescine dihydrochloride at a dose of 200 mg per kg body weight once daily for 7 consecutive days. Treatments were maintained throughout behavioral testing until tissue collection.

### Determination of Putrescine in Prefrontal Cortex

Following behavioral assessments, mice were euthanized under deep isoflurane anesthesia, and the prefrontal cortex (PFC) was rapidly dissected and weighed. Tissue homogenates were prepared by adding lysis buffer at a ratio of 500 µL per 0.1 g of tissue, followed by homogenization and incubation on ice for 10 minutes. Homogenates were centrifuged at 12,000 rpm for 5 minutes at 4°C, and the supernatant was collected for putrescine quantification.

Putrescine concentrations were measured using a commercial enzyme-linked immunosorbent assay (ELISA) kit (Enzyme-linked, REF: YJ523211) according to the manufacturer’s instructions. Samples were diluted 1:5 in dilution buffer, and 50 µL of each diluted sample or standard was added to pre-coated wells together with 100 µL of horseradish peroxidase (HRP)–conjugated detection antibody. After incubation for 60 minutes at 37°C, wells were washed five times, followed by the addition of substrate solution and incubation in the dark for 15 minutes. The reaction was terminated with stop solution, and absorbance was measured at 450 nm. Putrescine concentrations were calculated from a standard curve and corrected for the dilution factor.

### Immunoblotting

Prefrontal cortex tissues were homogenized individually in ice-cold lysis buffer supplemented with a protease inhibitor cocktail [48]. Protein concentrations were determined, and 20 µg of total protein per sample was loaded onto 10% SDS–polyacrylamide gels for electrophoretic separation, followed by transfer onto 0.2-µm polyvinylidene difluoride (PVDF) membranes using an electroblotting system.

The PVDF membranes were blocked in TBST blocking buffer for 16 hours at 4°C, followed by incubation overnight at 4°C with primary antibodies: anti-ATP13A3 (1:500, Proteintech, Cat# 55108-1-AP), anti-ATP13A2 (1:200, Santa Cruz, Cat# sc-293367), and anti-β-actin (1:10000, Santa Cruz, Cat# sc-47778).

Following three washes with TBST, the blots were incubated with goat anti-mouse IgG labeled with horseradish peroxidase (HRP) (1:1000, Beyotime, Cat# A0216) or goat anti-rabbit IgG labeled with HRP (1:1000, Beyotime, Cat# A0208) and visualized using a chemiluminescent HRP detection kit (Millipore, Cat# WBKLS0500). Images of the blots were captured using the ImageQuant LAS-4000 system, and the density of protein bands was quantified using ImageQuant TL software (version 7.0). The relative protein quantity was determined by normalizing the density of the target protein band to the density of the β-actin band in the same lane. Western blot analysis included two or six technical replicates for each sample.

### Fluorescent Labeling of PS128

Overnight cultures of *Lactiplantibacillus plantarum* PS128 were inoculated into sterile MRS broth at a 1% (v/v) ratio and grown to an optical density at 600 nm (OD_600_) of 1.0. Bacterial cells were harvested by centrifugation at 5,000 × g for 2 minutes at 4°C and washed three times with phosphate-buffered saline (PBS). Carboxyfluorescein diacetate succinimidyl ester (CFDA/SE, MedChemExpress, Cat# HY-D0938) was then added to a final concentration of 20 µM, and bacteria were incubated in the dark at 37°C for 30 minutes. Excess dye was removed by washing the cells three times with PBS.

Following labeling, PS128 cells were resuspended in PBS at concentrations of 1 × 10^9^ or 1 × 10^10^ colony-forming units (CFU) per milliliter. A small aliquot of the bacterial suspension was mounted onto microscope slides, and fluorescence images were acquired using a fluorescence microscope. Labeling efficiency was calculated as the proportion of fluorescently labeled cells relative to the total number of cells. Quantification was performed by randomly selecting seven fields of view from samples at 1 × 10^9^ CFU/mL and three fields of view from samples at 1 × 10^10^ CFU/mL.

### Determination of PS128 Growth Before and After CFDA/SE Treatment

PS128 cultures at a concentration of 1 × 10^10^ colony-forming units (CFU) per milliliter, with or without CFDA/SE labeling, were inoculated into sterile MRS broth at a 3% (v/v) ratio. Bacterial growth was monitored by measuring the optical density at 600 nm (OD_600_) every 30 minutes during the first 2 hours and then every 1 hour up to 8 hours to generate growth curves. For each condition, OD_600_ values at each time point were calculated as the mean of two technical replicates from five independent bacterial cultures.

### Determination of PS128 Tolerance to Gastric and Intestinal Fluids

Tolerance of *Lactiplantibacillus plantarum* PS128 to simulated gastric fluid (SGF, Phygene, Cat# PH1840) and simulated intestinal fluid (SIF, Phygene, Cat# PH1841) was assessed based on previously described methods by Corcoran et al. [49] and Azat et al. [50], with modifications. Briefly, PS128 cells resuspended in phosphate-buffered saline (PBS) were collected by centrifugation at 5,000 × g for 2 minutes at 4°C and resuspended in 2.5 mL of MRS broth for a recovery period of 2.5 hours.

An 800-µL aliquot of PS128 culture at a concentration of 1 × 10^10^ colony-forming units (CFU) per milliliter was mixed with an equal volume of MRS broth (control), a 1:1 mixture of MRS broth and SGF, or a 1:1 mixture of MRS broth and SIF. Cultures containing SGF were overlaid with mineral oil (Acmec, Cat# M97740) to maintain anaerobic conditions and incubated statically for 3 hours, whereas cultures containing SIF were incubated with shaking at 160 rpm for 4 hours. Cultures in MRS broth alone served as controls.

After incubation, 200 µL of each bacterial suspension at a 10^-5^ dilution was plated onto MRS agar plates. Following incubation at 37°C for 24 hours, colonies were imaged and counted using ImageJ software (version 1.54f). Tolerance was calculated as the percentage of colony-forming units recovered after SGF or SIF exposure relative to the MRS control. Six independent experiments were performed for statistical analysis.

### Detection and relative quantification of PS128 in fecal samples

Fresh fecal samples were collected from 3.5-month-old mice and incubated with lysozyme (3 mg/mL, Tiangen, Cat# RT401) in lysis buffer containing 10 mM Tris and 1 mM EDTA (pH 8.0, Sangon Biotech, Cat# EB0185) at 30°C for 5 minutes to facilitate bacterial cell lysis. Total RNA was isolated using TRIzol reagent according to the manufacturer’s instructions (TransGen Biotech, Cat# ER501). Complementary DNA (cDNA) was synthesized using the FastKing RT kit (Tiangen, Cat# KR116) following the manufacturer’s protocol. A positive control was prepared using 3 mL of PS128 culture at an optical density at OD_600_ of 0.6.

PS128-specific primers (Fig. S3V) for PCR and real-time reverse transcription quantitative PCR (RT-*q*PCR) [51] were designed based on the complete PS128 genome generated in this study and deposited in the NCBI database (BioProject accession number PRJNA1345344). PCR amplification was performed using 2× Taq Plus Master Mix II (Vazyme, Cat# P213-02) with an initial denaturation step at 94°C for 3 minutes, followed by 40 cycles of denaturation at 94°C for 30 seconds, annealing at 48°C for 35 seconds, and extension at 72°C for 25 seconds, with a final extension at 72°C for 10 minutes. Ten microliters of each PCR product were resolved on a 2% agarose gel and visualized under ultraviolet illumination using a gel documentation system.

RT-*q*PCR was performed using SYBR Green Master Mix (Qiagen, Cat# 208054) on a CFX Connect Real-Time PCR Detection System with 35 cycles consisting of denaturation at 95°C for 5 seconds followed by annealing and extension at 60°C for 30 seconds. Data were analyzed using Bio-Rad CFX Maestro software (version 3.1.1517.0823). Relative PS128 abundance was calculated using the 2^-ΔΔCt^ method, with bacterial 16S rRNA serving as the internal reference.

### Imaging the Localization of PS128 in the Mouse Gut

Adult mice in the PS128 group were administered 500 µL of fluorescently labeled PS128 (1 × 10^10^ CFU/mL) by oral gavage once daily for 3 consecutive days, whereas control mice received 500 µL of phosphate-buffered saline (PBS). After the final gavage, mice were divided into two groups for analysis.

For the first group, mice were anesthetized with isoflurane, and the abdominal area was shaved to facilitate in vivo fluorescence imaging using an IVIS Lumina III Smart Imaging System. Following whole-body imaging, mice were euthanized by cervical dislocation under deep isoflurane anesthesia, and the entire gastrointestinal tract was immediately isolated. Ex vivo fluorescence imaging of the gut was performed using the IVIS system with an excitation wavelength of 480 nm and an emission wavelength of 520 nm. Total and mean fluorescence intensities within defined regions of interest were quantified using Living Image software (version 4.5.2).

For the second group, mice were euthanized immediately after the final gavage. Segments of the gastrointestinal tract were collected, including a 10-cm segment of the small intestine proximal to the stomach, the cecum, and a 1.5-cm segment of the distal colon near the anus. Tissues were cut into approximately 5-mm squares and imaged using a stereoscopic fluorescence microscope. Mean fluorescence intensity was quantified using ImageJ software.

### Whole-genome sequencing and annotation of *Lactiplantibacillus plantarum* PS128

Whole-genome sequencing of *Lactiplantibacillus plantarum* PS128 was performed by BGI Genomics Co., Ltd. (Shenzhen, China) using a hybrid sequencing strategy. Detailed protocols for sequencing and annotation are provided in Supplementary Material 1. Whole-genome sequencing data for PS128 have been deposited in the NCBI database (BioProject accession number PRJNA1345344).

### 16S rRNA Gene Sequencing and analysis

DNA extraction, PCR amplification, library construction, and sequencing were performed by Origin-gene Biotechnology Co., Ltd. (Shanghai, China). Fecal samples were collected from four mouse groups (WT, WTPS, KO, and KOPS) at 8:00 p.m. Microbial genomic DNA was extracted using the QIAamp Fast DNA Stool Mini Kit (Qiagen, Germany) and assessed by 1% agarose gel electrophoresis. Comprehensive protocols for 16S rRNA gene sequencing and analysis are provided in Supplementary Material 1. The 16S rRNA gene sequencing data have been deposited in the NCBI Sequence Read Archive (BioProject accession number PRJNA1345244).

### Metagenomic Sequencing and Analysis

#### Sample preparation and sequencing

Total genomic DNA was extracted from 0.2 g of fecal sample using the MagAttract PowerSoil Pro DNA Kit (Qiagen, Germany) according to the manufacturer’s instructions. DNA concentration and purity were measured using a Quantus Fluorometer (PicoGreen, Promega, USA) and a NanoDrop 2000 spectrophotometer (Thermo Fisher Scientific, USA), respectively, and DNA integrity was assessed by 1% agarose gel electrophoresis.

Extracted DNA was fragmented to an average size of approximately 350 bp using a Covaris M220 Focused-ultrasonicator for paired-end library construction. Sequencing libraries were prepared using the NEXTFLEX Rapid DNA-Seq Kit (Bioo Scientific, USA) following the manufacturer’s protocol and sequenced on an Illumina NovaSeq 6000 platform (S4 flow cell). DNA extraction, library construction, and sequencing were performed by Majorbio Bio-Pharm Technology Co., Ltd. (Shanghai, China). Full methodological details for metagenomic sequencing and downstream analyses, including metagenomic assembly and annotation, reconstruction of metagenome-assembled genomes (MAGs), and gut–brain module (GBM) analysis, are described in Supplementary Material 1. Metagenomic sequencing data have been deposited in the NCBI Sequence Read Archive (BioProject accession number PRJNA1345245).

### Single-nucleus RNA sequencing

#### Frozen Tissue Nuclear Dissociation

Single-nucleus RNA sequencing (snRNA-seq) was performed by Shanghai Majorbio Bio-Pharm Technology Co., Ltd. (Shanghai, China). Frozen prefrontal cortex (PFC) tissues were homogenized using a glass Dounce tissue grinder (Sigma, USA) with 25 strokes of pestle A and 25 strokes of pestle B in 2 mL of ice-cold Nuclei EZ lysis buffer. The homogenate was incubated on ice for 5 minutes, followed by addition of 3 mL of cold lysis buffer and an additional 5-minute incubation. Nuclei were centrifuged at 500 × g for 5 minutes at 4°C, washed with 5 mL of ice-cold lysis buffer, and incubated on ice for an additional 5 minutes. The nuclear pellet was resuspended and washed in 5 mL of nuclei suspension buffer (NSB; PBS supplemented with 0.01% BSA and 0.1% RNase inhibitor, Clontech, USA). Nuclei were resuspended in 2 mL of NSB, filtered through a 35-µm cell strainer (Corning-Falcon, USA), and counted. Nuclei were loaded onto the 10x Genomics Chromium Controller at a final concentration of 1,000 nuclei per µL.

#### 10x library preparation and sequencing

Gel beads containing unique molecular identifiers (UMIs) and cell barcodes were loaded to near saturation such that each nucleus was paired with one bead in a Gel Bead-in-Emulsion (GEM). After lysis, polyadenylated RNA molecules hybridized to bead-bound oligonucleotides. Beads were collected for reverse transcription, during which each cDNA molecule was tagged at its 3’ end (corresponding to the 5’ end of the mRNA) with a UMI and cell barcode. Subsequent steps included second-strand synthesis, adaptor ligation, and universal amplification. Sequencing libraries were prepared from randomly fragmented cDNA to enrich 3’ transcript ends linked to barcodes and UMIs.

All procedures followed the Chromium Single Cell 3’ GEM, Library & Gel Bead Kit (v3.1 chemistry) manufacturer’s protocol. Libraries were quantified using a High Sensitivity DNA Chip (Agilent) on a Bioanalyzer 2100 and a Qubit High Sensitivity DNA Assay (Thermo Fisher Scientific), and sequenced on an Illumina NovaSeq X Plus using paired-end 150-bp chemistry. Detailed methods for snRNA-seq data processing, differential expression, and functional enrichment analyses are provided in Supplementary Material 1. Results of snRNA-seq have been deposited in the NCBI Sequence Read Archive (BioProject accession number PRJNA1345209).

### Targeted Metabolomics of Fatty Acids and Neurotransmitters in Serum and Brain

#### Targeted metabolomics for fatty acids in brain

Mice were anesthetized with isoflurane, after which venous blood was collected by ophthalmectomy. Brain tissues were harvested, immediately frozen in liquid nitrogen, and stored at −80°C until analysis. Serum was prepared by sequential centrifugation and stored at −80°C until use. Targeted metabolomics analyses were performed by Shanghai BioTree Biomedical Technology Co., Ltd. Brain lipid extraction, sample preparation, and GC–MS–based fatty acid quantification were conducted using standardized protocols. Detailed procedures, including tissue processing, solvent extraction, instrument parameters, and data acquisition settings, are provided in Supplementary Material 1.

#### Targeted metabolomics for neurotransmitters in serum and brain

Targeted metabolomics of neurotransmitters in serum and brain was performed using UHPLC–MS/MS following established extraction, derivatization, and analytical procedures. Detailed sample preparation protocols, chromatographic conditions, mass spectrometry parameters, and data processing workflows are provided in Supplementary Material 1.

### Quantification and statistical analysis

All behavioral and histological analyses were performed under blinded conditions. Statistical analyses were carried out using GraphPad Prism (GraphPad Software, version 10.6.1). Normality was assessed using the Shapiro–Wilk, D’Agostino–Pearson, Anderson–Darling, and Kolmogorov–Smirnov tests. Data are presented as mean ± SEM unless otherwise stated. Normally distributed data were compared using two-tailed (or one-tailed, as specified in figure legends) unpaired Student’s *t*-tests. Non-normally distributed data were analyzed using two-tailed (or one-tailed, as specified) Mann–Whitney U tests. *P* values < 0.05 were considered statistically significant. Exact sample sizes (n) and statistical tests used are reported in the figure legends, and precise *P* values are indicated in the figures. All raw data used in this study are provided in Supplementary Tables S1 and S2.

## Supplemental information

**Supplementary Material 1.**

Supplementary Methods and Figs. S1–S7 related to Figs. 1–6.

**Supplementary Material 2.**

Table S1. Raw data related to Figs. 1–6.

**Supplementary Material 3.**

Table S2. Raw data related to Figs. S1–S7.

**Supplementary Material 4.**

Video S1. Voluntary intake of PS128-containing or blank jelly by mice.

**Supplementary Material 5.**

Video S2. Behavioral statistics from the olfactory habituation/dishabituation test, related to Fig. S1.

**Supplementary Material 6.**

Video S3. Four specific behaviors in the olfactory habituation/dishabituation test, related to Fig. S1.

## Supporting information

Raw data related to Figures 1_6

Raw data related to Figures S1_S7

Four specific behaviors in OHDT, related to Fig. S1

Behavioral data statistics from OHDT, related to Fig. S1

Voluntary intake of probiotics-containing and blank jelly

## Acknowledgements

We thank the ECNU Multifunctional Platform for Innovation (101) for their mouse breeding service.

## Authors’ contributions

YHP and XBY conceived and supervised the project and obtained funding. YMC, YXC, and SZ conducted behavioral, histological, and molecular analyses. JLC, YXC, YMC, TAL, and SSG conducted data processing. YMC, YXC, and YHP conducted data analysis and figure organization. YHP, XBY, YCT, YWL, JM, WZ, and HPL contributed to the discussion and data interpretation. YMC, YXC, YHP, and XBY wrote the manuscript. All authors read and approved the final manuscript.

## Data availability

Whole-genome sequencing data for *Lactiplantibacillus plantarum* PS128 have been deposited in NCBI-SRA under accession number PRJNA1345344. 16S rRNA sequencing data have been deposited in NCBI-SRA under accession number PRJNA1345244. Metagenomic sequencing data have been deposited in NCBI-SRA under accession number PRJNA1345245. Single-nucleus RNA sequencing data have been deposited in NCBI-SRA under accession number PRJNA1345209.

## Declarations

### Ethics approval and consent to participate

All animal procedures were approved by the Institutional Animal Care and Use Committee of East China Normal University (approval no. m20241106) and were conducted in accordance with institutional guidelines.

### Consent for publication

Not applicable.

### Author details

^1^Key Laboratory of Brain Functional Genomics of Shanghai and Ministry of Education, Institute of Brain Functional Genomics, School of Life Science and the Collaborative Innovation Center for Brain Science, East China Normal University, Shanghai 200062, China. ^2^Institute of Biochemistry and Molecular Biology, National Yang Ming Chiao Tung University, Taipei 11221, Taiwan. ^3^Institute of Mechanobiology and Medical Engineering, School of Life Sciences and Biotechnology, Shanghai Jiao Tong University, Shanghai 201100, China. ^4^Shanghai Institute of Nutrition and Health, University of Chinese Academy of Sciences, Chinese Academy of Sciences, Shanghai 200031, China.

