## Supplementary Material 1 for "*Lactiplantibacillus plantarum* PS128 restores microbiota-driven polyamine homeostasis to alleviate autism-like behaviors in *Fmr1* knockout mice"

[1]Key Laboratory of Brain Functional Genomics of Shanghai and Ministry of Education, Institute of Brain Functional Genomics, School of Life Science and the Collaborative Innovation Center for Brain Science, East China Normal University, Shanghai 200062, China
[2]Institute of Biochemistry and Molecular Biology, National Yang Ming Chiao Tung University, Taipei 11221, Taiwan
[3]Institute of Mechanobiology and Medical Engineering, School of Life Sciences and Biotechnology, Shanghai Jiao Tong University, Shanghai 201100, China
[4]Shanghai Institute of Nutrition and Health, University of Chinese Academy of Sciences, Chinese Academy of Sciences, Shanghai 200031, China.
[5]These authors contributed equally
[6]Lead contact

**Supplementary Material 1.**
Supplementary Methods and Figs. S1–S7 related to Figs. 1–6.
**Supplementary Material 2.**
Table S1. Raw data related to Figs. 1–6.
**Supplementary Material 3.**
Table S2. Raw data related to Figs. S1–S7.
**Supplementary Material 4.**
Video S1. Voluntary intake of PS128-containing or blank jelly by mice.
**Supplementary Material 5.**
Video S2. Behavioral statistics from the olfactory habituation/dishabituation test, related to Fig. S1.
**Supplementary Material 6.**
Video S3. Four specific behaviors in the olfactory habituation/dishabituation test, related to Fig. S1.

### Methods

#### Behavioral Experiments

***The open-field test***

The open-field test was performed in a square arena with white opaque walls (40 cm × 40 cm × 40 cm). Mice were gently placed in the center of the arena and allowed to explore individually for 5 minutes under an illumination of 45 lux. Total distance traveled and time spent in the center zone (20 cm × 20 cm) were quantified.

***Novel object recognition test***

The test was performed in the same arena used for the open-field test under an illumination of 3 to 5 lux. The test consisted of two 10-minute sessions separated by a 1-hour interval. During the first session (learning phase), mice were allowed to freely explore the arena containing two identical objects (Object A1 and Object A2; 6 × 6 × 10 cm, length × width × height). After the 1-hour interval, mice were reintroduced to the arena for the second session (testing phase), during which one familiar object was replaced with a novel object of identical dimensions (Object B). Sniffing behavior was defined as the mouse directing its nose toward an object at a distance within 3 cm. The total duration and frequency of sniffing the novel object were quantified and compared with those for the familiar object. Climbing behavior was quantified as the total duration and frequency of climbing onto Objects A and B and was defined as any event in which both hind limbs were off the ground. The recognition index was calculated as the percentage of time spent sniffing the novel object relative to the total sniffing time.

***Elevated plus maze test***

The elevated plus maze consisted of a plus-shaped platform with two enclosed arms (30 cm × 5 cm) surrounded by 15 cm-high walls and two open arms (30 cm × 5 cm). The maze was elevated 60 cm above the floor. At the start of the test, mice were placed in the central area of the maze facing an open arm and allowed to explore freely for 5 minutes under an illumination of 45 lux. The time spent and distance traveled in the open and enclosed arms were quantified.

***The O-maze test***

The O-maze test was performed using a circular elevated track constructed of gray acrylic, with a height of 40 cm, an inner diameter of 48 cm, and a width of 5 cm. The maze consisted of two open arms and two enclosed arms of equal length arranged alternately along the track, with each arm spanning 45°. The open arms were equipped with 0.5 cm-high acrylic strips serving as anti-fall guardrails, whereas the enclosed arms were surrounded by 15 cm-high walls to block visual access to the external environment.

At the start of the test, each mouse was placed facing an enclosed arm and allowed to explore freely for 5 minutes under an illumination of 45 lux. Time spent, distance traveled, and the number of entries into the open and enclosed arms were quantified.

***Balance beam test***

The test was performed using an inclined square wooden beam with a flat surface. For the 30° condition, the beam was 0.8 cm wide and 100 cm long, with the lower and higher ends positioned at vertical heights of 26 cm and 76 cm, respectively. A black shelter box (12 × 12 × 12 cm; wall thickness, 3 mm) containing nesting material previously used by the mice for 3 days was placed at the higher end of the beam.

During the training phase, mice were placed at the lower end of the 30° beam and guided to traverse the beam once per day for 3 consecutive days. If a mouse paused for more than 10 seconds without forward movement, it was gently encouraged to proceed using the back of the experimenter's index finger. During the testing phase, mice traversed the beam without guidance and were allowed to rest in the shelter box for 15 seconds after crossing. Each mouse completed four trials on the 30° beam. The beam was then replaced with a narrower beam (0.6 cm wide and 50 cm long), with the lower and higher ends positioned at 33 cm and 76 cm, respectively, resulting in a 60° incline. Mice completed one trial on the 60° beam, and the latency to traverse the beam was recorded.

Performance was scored on a 7-point scale. A full score of 7 was assigned when a mouse crossed the beam without slipping, falling, or requiring prompting. One point was deducted for each occurrence of slipping, falling, or prompting, with a minimum score of 1.

***Reciprocal social interaction test***

The test was performed in the same arena used for the open-field test. The test consisted of three 3-minute sessions separated by 30-minute rest intervals. In each session, the experimental mouse was placed in the arena and allowed to interact with an age-, sex-, and weight-matched conspecific.

During the first session, the experimental mouse interacted with an unfamiliar wild-type (WT) mouse (Mouse 1). After a 30-minute rest interval, the experimental mouse was reintroduced to the arena to interact with the same WT mouse (Mouse 1), which had become familiar, during the second session. Following a second 30-minute rest interval, the experimental mouse interacted with a different unfamiliar WT mouse (Mouse 2) during the third session. Social behaviors, including chasing, physical contact, sniffing within a 2-cm interaction zone, and crawling over WT mouse were quantified.

***Olfactory habituation/dishabituation test***

The olfactory habituation/dishabituation test was performed in a clean, transparent cage (27.2 × 17.5 × 11.5 cm, length × width × height) lined with fresh bedding and covered with a lid containing a 5-mm-diameter hole for odor presentation. Each mouse was placed in the cage and allowed to acclimate for 90 seconds before testing.

During testing, a cotton swab saturated with 100 µL of a given odor was introduced through the hole and suspended in the cage for 1 minute. The swab was replaced every minute with a fresh one, resulting in three consecutive presentations per odor. This procedure was repeated sequentially for four odors: pure water, imitation banana flavor (1:100 dilution, McCormick & Co., Inc.), and soiled bedding collected from two independent cages, each housing two sibling wild-type (WT) mice.

Water and banana flavor were used as nonsocial odors. Soiled bedding from sex- and age-matched WT mice was used as a social odor. Social odor samples were obtained from two distinct cages of sibling WT mice that had been co-housed for at least 7 days, ensuring different family lineages between cages. Odors were collected by rubbing a cotton swab along the bottom surface of each cage in a circular motion six times.

The duration of sniffing, nibbling, reaching toward, and maintaining contact with the swab was quantified, and the proportion of each behavior relative to the total interaction time was calculated (Videos S2 and S3).

***Tube test***

The tube test was performed using a transparent acrylic tube (30 cm in length, 3 cm inner diameter, 3 mm wall thickness). To facilitate habituation, two identical 15 cm tubes were placed in the home cages for three consecutive days. Mice were then trained to traverse the 30 cm tube six times per day (three passes in each direction) for three consecutive days.

On the testing day, weight-matched mice from different experimental groups (WT vs. KO, WT vs. WTPS, WT vs. KOPS, or KO vs. KOPS) were placed at opposite ends of a horizontally positioned tube and allowed to enter simultaneously. The mouse that advanced through the tube was assigned a score of 1, whereas the mouse that retreated was assigned a score of 0. After an interval of 2 hours, a second trial was conducted, pairing mice with similar first-trial scores and body weights.

**Whole-genome sequencing and annotation of *Lactiplantibacillus plantarum* PS128**

Long-read sequencing was conducted on the PacBio RS II platform, and short-read sequencing was performed on the Illumina HiSeq 4000 platform. PacBio subreads shorter than 1 kb were removed during quality filtering. Error correction was performed using Pbdagcon, and corrected reads were assembled into draft contigs using the Celera Assembler. Illumina short reads were subsequently used to polish the assembly and improve base accuracy.

Gene prediction was performed using Glimmer3. Transfer RNAs, ribosomal RNAs, and small RNAs were identified using standard RNA annotation tools. Tandem repeats, genomic islands, prophage regions, and CRISPR arrays were identified using established bioinformatics pipelines.

Functional annotation of predicted genes was conducted by sequence similarity searches against public databases, including KEGG, COG, NR, Swiss-Prot, GO, TrEMBL, and EggNOG. Genes associated with virulence, antimicrobial resistance, host–pathogen interaction, and carbohydrate metabolism were annotated using specialized databases.

Comparative genomic analyses were performed against representative *Lactiplantibacillus* genomes using whole-genome alignment and clustering approaches. Core and pan-genome analyses were conducted using CD-HIT, and phylogenetic relationships were inferred based on whole-genome comparisons.

**16S rRNA Gene Sequencing and analysis**

The V3–V4 region of the bacterial 16S rRNA gene was amplified by PCR using primers 338F (5’-ACTCCTACGGGAGGCAGCAG-3’) and 806R (5’-GGACTACHVGGGTWTCTAAT-3’), each containing an eight-base sample-specific barcode. PCR was performed with an initial denaturation at 95°C for 3 minutes, followed by 27 cycles of denaturation at 95°C for 30 seconds, annealing at 55°C for 30 seconds, and extension at 72°C for 45 seconds, with a final extension at 72°C for 10 minutes. Reactions were carried out in triplicate 20-μL volumes containing 5× FastPfu Buffer, dNTPs, primers, FastPfu Polymerase, and 10 ng of template DNA. Amplicons were excised from 2% agarose gels and purified using the AxyPrep DNA Gel Extraction Kit (Axygen Biosciences, USA).

Sequencing libraries were prepared using the NEBNext Ultra DNA Library Prep Kit for Illumina (NEB, USA) according to the manufacturer's instructions. Library quality was assessed using a Qubit 2.0 Fluorometer (Thermo Scientific, USA) and an Agilent Bioanalyzer 2100 system. Paired-end sequencing (PE250) was performed on an Illumina NovaSeq 6000 platform.

Raw sequencing data were processed using VSEARCH (version 2.15.2) on a Linux platform [1]. After removal of primers and low-quality reads, sequences were merged, dereplicated, and clustered into operational taxonomic units (OTUs) at 97% sequence similarity, with chimeric sequences removed based on the RDP Gold database. An OTU abundance table was generated, and taxonomic classification was assigned using the SILVA reference database (release 126).

Downstream analyses were performed in R. The OTU table and taxonomic annotations were imported into a phyloseq object. Alpha-diversity metrics, including observed species, Shannon index, and Simpson index, were calculated using the microbiome package. Beta-diversity analyses were conducted based on Bray–Curtis and UniFrac distances, followed by principal coordinates analysis using the vegan package. Differentially abundant taxa were identified using LEfSe analysis implemented on the OmicStudio platform, and data visualization was performed using ggplot2.

**Metagenomic Sequencing and Analysis**

***Metagenomic assembly and annotation***

Raw sequencing reads were quality-filtered using FASTP (version 0.23.0) to remove low-quality reads and reads containing ambiguous bases [2]. Host-derived reads were removed by alignment to the mouse reference genome using BWA (version 0.7.17) [3]. High-quality reads were assembled de novo using MEGAHIT (version 1.1.2) [4], and contigs ≥300 bp were retained. Open reading frames were predicted using Prodigal (version 2.6.3) [5], and a non-redundant gene catalog was generated using CD-HIT (version 4.7) [6].

Gene abundance was estimated by mapping quality-filtered reads back to the gene catalog using SOAPaligner (version 2.21) [7]. Taxonomic annotation was performed by aligning non-redundant genes to the NCBI non-redundant (NR) database using DIAMOND (version 2.0.13) [8]. Functional annotation was conducted based on the Kyoto Encyclopedia of Genes and Genomes (KEGG, www.kegg.jp) database. Data visualization and statistical analyses were performed in R using ggplot2.

***Reconstruction of metagenome-assembled genome (MAG) and gut-brain module (GBM) analysis***

After quality control, removal of host-derived reads, and de novo assembly, contigs were binned into metagenome-assembled genomes (MAGs) using MetaBAT2 (version 2.15) based on sequence composition and coverage depth, yielding 1,297 bins. MAG quality was assessed using CheckM (version 1.2.2) to estimate genome completeness and contamination. High-quality MAGs were defined as those with ≥ 50% completeness and ≤ 10% contamination; bins not meeting these thresholds were classified as low quality. Using these criteria, 533 high-quality MAGs were retained.

To remove redundancy, high-quality MAGs were dereplicated using dRep (version 3.4.2), resulting in 200 unique, non-redundant MAGs. Taxonomic assignment was performed using DIAMOND against the NCBI Taxonomy database to define species-level genome bins (SGBs). Protein-coding genes within MAGs were annotated using DIAMOND against the NCBI non-redundant (NR) protein database. Functional annotation and enrichment analyses were performed using KofamKOALA (version 1.3.0) with the Kyoto Encyclopedia of Genes and Genomes (KEGG) database.

Gut–brain modules (GBMs) encoded by SGBs were reconstructed based on curated definitions described by Valles-Colomer et al. [9], and categorized into eight functional subgroups according to their roles. Visualization and plotting were performed in R using ggplot2 and Chiplot.

### Single-nucleus RNA sequencing

#### *snRNA-seq data processing*

Raw reads were processed using Cell Ranger (version 7.1.0) with default parameters. FASTQ files generated from Illumina sequencing were aligned to the mouse GRCm39 genome (Ensembl release 111 annotation) using the STAR algorithm. Gene–barcode matrices were generated by counting UMIs and filtering non-cell-associated barcodes. The resulting matrix was imported into Seurat (version 4.4.3) for quality control and downstream analysis [10]. Low-abundance genes were filtered such that only features expressed in at least three cells were retained. Cells were filtered using the following criteria: 200 to 6,000 detected genes per cell and mitochondrial gene percentage < 25% (PercentageFeatureSet). Data were normalized using NormalizeData, and the top 2,000 highly variable genes were identified using FindVariableFeatures. Principal component analysis (PCA) was performed, and 12 significant principal components (PCs) were selected based on the ElbowPlot function. A shared nearest-neighbor (SNN) graph was constructed using FindNeighbors, and clusters were identified using FindClusters at a resolution of 0.3, optimized with Clustree (version 0.5.1) [11]. Clusters were visualized in Uniform Manifold Approximation and Projection (UMAP) space.

Differentially expressed genes (DEGs) were identified using the Wilcoxon rank-sum test (FindAllMarkers). Cell types were annotated using a semi-automated strategy based on the CellMarker 2.0 database [12] and published references by Tiklová et al. [13], Bhattacherjee et al. [14], Liu et al. [15], and Hing et al [16]. Expression of *Atp13a2*, *Atp13a3*, and *Atp13a4* was visualized using FeaturePlot_scCustom in scCustomize (version 3.2.0) and ggplot2 (version 3.5.2).

#### *Differential expression analysis and functional enrichment*

DEGs were identified using FindMarkers with the following thresholds: genes expressed in ≥ 10% of cells, adjusted $P \leq 0.05$, and $|\log_2 FC| > 0.12$. VennDiagram (version 1.7.3) was used to identify genes significantly upregulated or downregulated in KOPS vs. KO (adjusted $P \leq 0.05$ and $|\log_2 FC| > 0.12$) but not significantly different from WT (adjusted $P > 0.05$).

Gene Ontology (GO) enrichment analysis was performed using enrichGO in clusterProfiler (version 4.14.6) after gene ID conversion with org.Mm.eg.db (version 3.20.0) [17]. Enriched

Biological Process (BP) terms were defined at $q \leq 0.05$ (Benjamini–Hochberg adjusted) using annotations from the JAX Gene Ontology Database and visualized using ggplot2 (version 3.5.2) [18].

**Targeted Metabolomics of Fatty Acids and Neurotransmitters in Serum and Brain**

***Targeted metabolomics for fatty acids in brain***

Mice were anesthetized with isoflurane, after which venous blood was collected by ophthalmectomy. Brain tissues were then harvested, immediately frozen in liquid nitrogen, and stored at −80°C until analysis. Blood samples were allowed to stand at room temperature for 1 hour before centrifugation at 3,000 rpm for 10 minutes. The supernatant was centrifuged again at 12,000 rpm for 10 minutes at 4°C, and the resulting serum was collected and stored at −80°C until use. All targeted metabolomics analyses were performed by Shanghai BioTree Biomedical Technology Co., Ltd.

Brain tissues were transferred to 2-mL Eppendorf tubes and extracted with 1 mL of $H_2O$, followed by vortex mixing for 10 seconds. Samples were homogenized in a ball mill at 40 Hz for 4 minutes and sonicated for 5 minutes in an ice-water bath. Homogenization and sonication were repeated three times. Samples were centrifuged at 5,000 rpm for 20 minutes at 4ºC, and 0.8 mL of the supernatant was transferred to a new 2-mL Eppendorf tube. To each tube, 0.1 mL of 50% $H_2SO_4$ and 0.8 mL of extraction solution (25 mg/L internal standard in methyl tert-butyl ether) were added. Samples were vortexed for 10 seconds, oscillated for 10 minutes, sonicated for an additional 10 minutes in an ice-water bath, and centrifuged at 10,000 rpm for 15 minutes at 4ºC. The supernatant was kept at −20ºC for 30 minutes and then transferred to 2-mL glass vials for GC–MS analysis.

Quantification of fatty acids was performed using a SHIMADZU GC2030-QP2020 NX gas chromatograph–mass spectrometer equipped with an HP-FFAP capillary column. A 1-μL aliquot of each extract was injected in split mode (5:1). Helium was used as the carrier gas at a flow rate of 1.2 mL/minute, with a front inlet purge flow of 3 mL/minute. The oven temperature program was as follows: 50ºC for 1 minute; ramp to 150ºC at 50ºC/minute (1-minute hold); ramp to 170ºC at 10ºC/minute; ramp to 225ºC at 20ºC/minute (1-minute hold); and ramp to 240ºC at 40ºC/minute (1-minute hold). The injection port, transfer line, quadrupole, and ion source temperatures were set to 240ºC, 240ºC, 150ºC, and 200ºC, respectively. Electron impact ionization was performed at −70 eV, and spectra were acquired in Scan/SIM mode over an m/z range of 33–150 following a 3.75-minute solvent delay.

***Targeted metabolomics for neurotransmitters in serum and brain***

For serum samples collected from isoflurane-anesthetized mice, 20 μL of serum was transferred to a 1.5-mL Eppendorf tube and mixed with 80 μL of precooled (−20ºC) extraction solvent (acetonitrile containing 0.1% formic acid). Samples were vortexed for 30 seconds, sonicated for 15 minutes in an ice-water bath, incubated at −40°C overnight, and centrifuged at 12,000 rpm for 15 minutes at 4ºC.

For brain samples, tissues were weighed and transferred to 1.5-mL Eppendorf tubes. Eighty microliters of precooled extraction solvent (acetonitrile containing 0.1% formic acid) and 20 μL of $H_2O$ were added, followed by vortexing for 30 seconds, homogenization at 45 Hz for 4 minutes, and sonication for 5 minutes in an ice-water bath. Homogenization and sonication were repeated three times. Samples were incubated at −20ºC overnight and centrifuged at 12,000 rpm for 15 minutes at 4ºC. Eighty microliters of each supernatant were mixed with 40 μL of 100 mM sodium carbonate and 40 μL of 2% benzoyl chloride in acetonitrile. After incubation for 30 minutes, 10 μL of internal standard was added, and samples were centrifuged at 12,000 rpm for 15 minutes at 4ºC. Forty microliters of the supernatant were diluted with 20 μL of $H_2O$ and transferred to autosampler vials for UHPLC–MS/MS analysis.

UHPLC separation was performed on an ACQUITY Premier System (Waters) equipped with an ACQUITY UPLC HSS T3 column (100 × 2.1 mm, 1.8 μm). The mobile phases were A: 0.1% formic acid and 1 mM ammonium acetate in water, and B: acetonitrile. The column temperature was maintained at 40ºC, the autosampler temperature at 10ºC, and the injection volume was 1 μL. Mass spectrometric detection was conducted on a SCIEX Triple Quad™ 6500+ mass spectrometer with the following parameters: IonSpray Voltage, +5,000 V; Curtain Gas, 35 psi; Source Temperature, 400ºC; Ion Source Gas 1 and 2, 60 psi. Data acquisition and processing were performed using SCIEX Analyst Workstation (version 1.6.3) and Data Driven Flow (version 2.0.3.11).

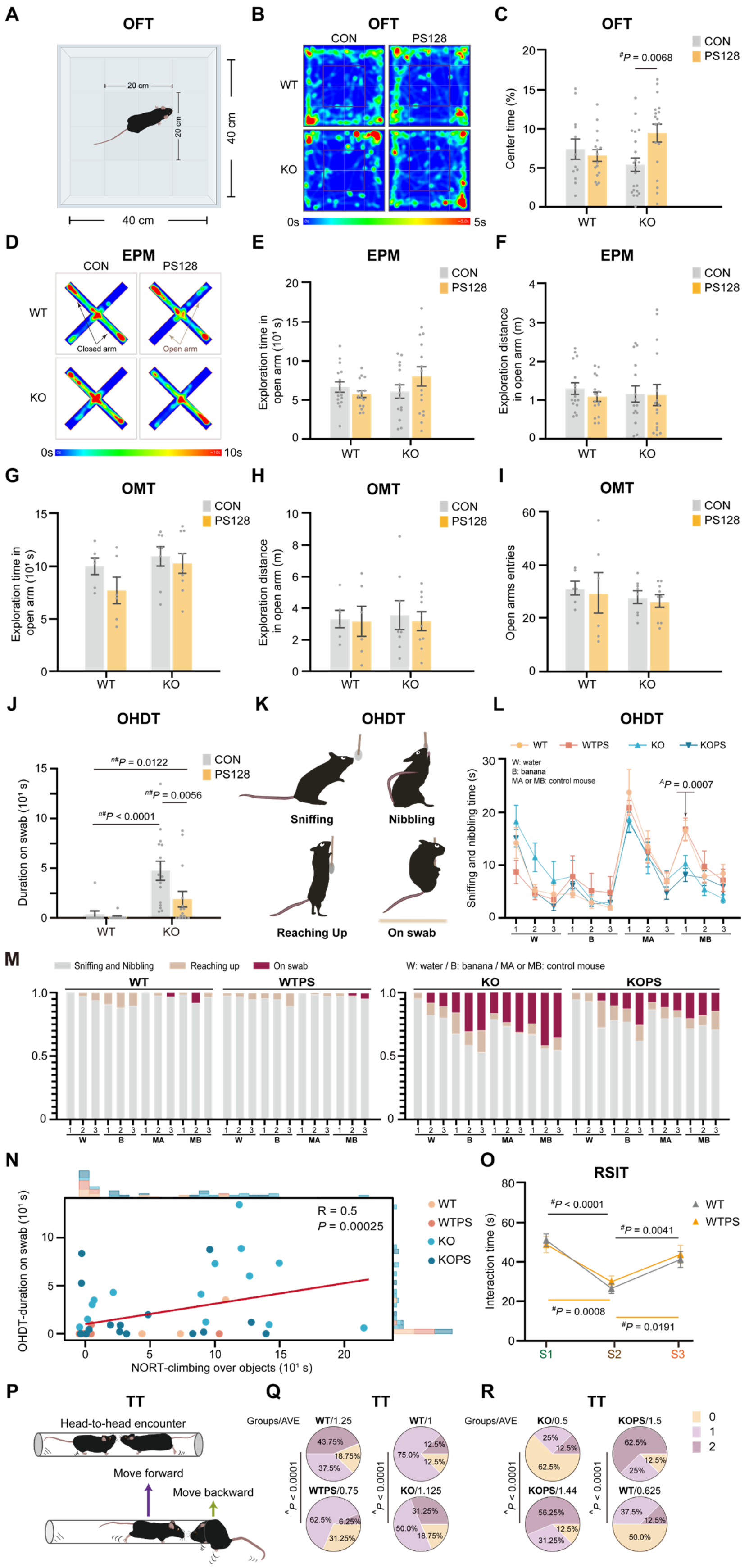

A
OFT
20 cm
20 cm
40 cm
40 cm
B
OFT
CON
PS128
WT
KO
0s
5s
C
OFT
CON
PS128
$^{\#}P$ = 0.0068
Center time (%)
WT
KO
D
EPM
CON
PS128
WT
KO
Closed arm
Open arm
0s
10s
E
EPM
CON
PS128
Exploration time in open arm (10¹ s)
WT
KO
F
EPM
CON
PS128
Exploration distance in open arm (m)
WT
KO
G
OMT
CON
PS128
Exploration time in open arm (10¹ s)
WT
KO
H
OMT
CON
PS128
Exploration distance in open arm (m)
WT
KO
I
OMT
CON
PS128
Open arms entries
WT
KO
J
OHDT
CON
PS128
$^{n\#}P$ = 0.0122
$^{n\#}P$ = 0.0056
$^{n\#}P$ < 0.0001
Duration on swab (10¹ s)
WT
KO
K
OHDT
Sniffing
Nibbling
Reaching Up
On swab
L
OHDT
WT
WTPS
KO
KOPS
W: water
B: banana
MA or MB: control mouse
$^{\Delta}P$ = 0.0007
Sniffing and nibbling time (s)
W
B
MA
MB
M
Sniffing and Nibbling
Reaching up
On swab
W: water / B: banana / MA or MB: control mouse
WT
WTPS
KO
KOPS
W
B
MA
MB
N
R = 0.5
P = 0.00025
WT
WTPS
KO
KOPS
OHDT-duration on swab (10¹ s)
NORT-climbing over objects (10¹ s)
O
RSIT
$^{\#}P$ < 0.0001
$^{\#}P$ = 0.0041
$^{\#}P$ = 0.0008
$^{\#}P$ = 0.0191
WT
WTPS
Interaction time (s)
S1
S2
S3
P
TT
Head-to-head encounter
Move forward
Move backward
Q
TT
Groups/AVE
WT/1.25
WT/1
WTPS/0.75
KO/1.125
$^{\Delta}P$ < 0.0001
43.75%
18.75%
37.5%
12.5%
75.0%
12.5%
62.5%
6.25%
31.25%
31.25%
50.0%
18.75%
R
TT
Groups/AVE
KO/0.5
KOPS/1.5
KOPS/1.44
WT/0.625
$^{\Delta}P$ < 0.0001
25%
12.5%
62.5%
62.5%
12.5%
25%
56.25%
12.5%
31.25%
37.5%
12.5%
50.0%
0
1
2

**Fig. S1 Behavioral characterization of *Fmr1* KO and wild-type mice across experimental groups, related to Fig. 1. A–C** Schematic of the open-field test (OFT) (**A**), heatmap of total time spent in the OFT (**B**), and time spent in the center area of the OFT (**C**), *n* = 12–21. **D–F** Heatmap of total time spent in the elevated plus maze (EPM) (**D**), exploration time (**E**) and distance (**F**) in the open arms of the EPM, *n* = 15–16. **G–I** Exploration time (**G**), distance traveled (**H**), and number of entries (**I**) in the open arms of the O-maze test, *n* = 6–9. **J–N** Total climbing time in each mouse group during the olfactory habituation/dishabituation test (OHDT) (**J**), schematic of animal postures (**K**), total time spent in sniffing and nibbling (**L**), proportions of sniffing/nibbling, reaching up, and climbing behaviors per mouse (**M**), and Spearman's rank correlation analysis between OHDT swab interaction duration and NORT climbing time (**N**), *n* = 10–18. W, water; B, banana; MA, control mouse A; MB, control mouse B. **O** Interaction time with familiar and novel mice across three stages in wild-type (WT) and PS128-supplemented WT (WTPS) mice in the reciprocal social interaction test (RSIT), *n* = 15–16. **P–R** Schematic of the tube test (TT) (**P**) and average score (AVE) per mouse obtained in the TT (**Q** and **R**), *n* = 16. Data are presented as mean ± S.E.M., and gray dots indicate individual mice. Two-tailed ($^{\#}$) Student's *t*-test and two-tailed ($^{n\#}$) Mann–Whitney U test were used for two-group comparisons; one-way ANOVA ($^{A}$) with post hoc Holm–Sidak tests was applied for multiple group comparisons. A chi-square (^) test was used in (**Q** and **R**). A *P* value < 0.05 was considered statistically significant.

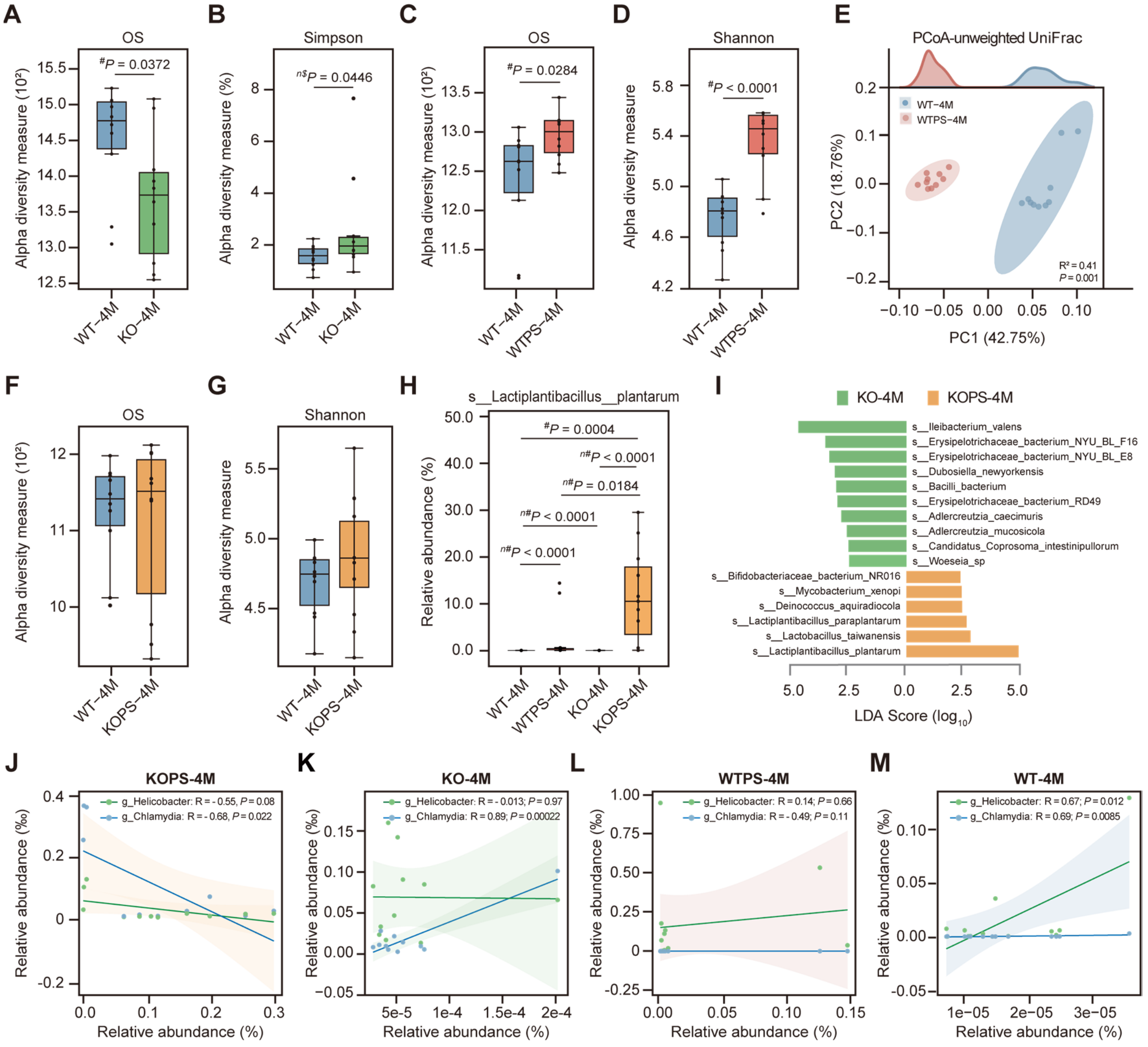


**Fig. S2 Gut microbiota diversity and composition across experimental mouse groups, related to Fig. 2. A** and **B** Gut microbiota analysis of 4-month-old *Fmr1* KO (KO-4M) and wild-type (WT-4M) mice without PS128 intervention. Observed species (**A**). Simpson's diversity index (**B**), $n$ = 10. **C-E** Gut microbiota analysis of WT-4M and PS128-supplemented WT (WTPS-4M) mice. Observed species (**C**). Shannon diversity index (**D**), and principal coordinates analysis (PCoA) plot (**E**). $n$ = 10. **F** and **G** Gut microbiota analysis of PS128-treated *Fmr1* KO (KOPS-4M) and WT-4M mice. Observed species (**F**). Shannon diversity index (**G**), $n$ = 10. **H** Elevated abundance of *Lactiplantibacillus plantarum* in KOPS-4M mice, $n$ = 11–13. **I** Gut microbiota analysis comparing 4-month-old *Fmr1* KO mice with or without PS128 treatment. Histogram of logarithmic linear discriminant analysis (LDA) scores, cutoff > 2.3, $P < 0.05$ (**I**), $n$ = 11. **J-M** Pearson correlation analyses between the relative abundances of *Lactiplantibacillus plantarum* and *Helicobacter* or *Chlamydia* in 4-month-old KOPS (**J**), KO (**K**), WTPS (**L**), and WT (**M**) mouse groups, $n$ = 11–13. Data are presented as mean ± S.E.M., and each dot represents individual mouse. Statistical significance was determined by two-tailed Student's $t$-test ($^{\#}$), one-tailed ($^{n\$}$) or two-tailed ($^{n\#}$) Mann–Whitney U test, as indicated. A $P$ value < 0.05 was considered significant.

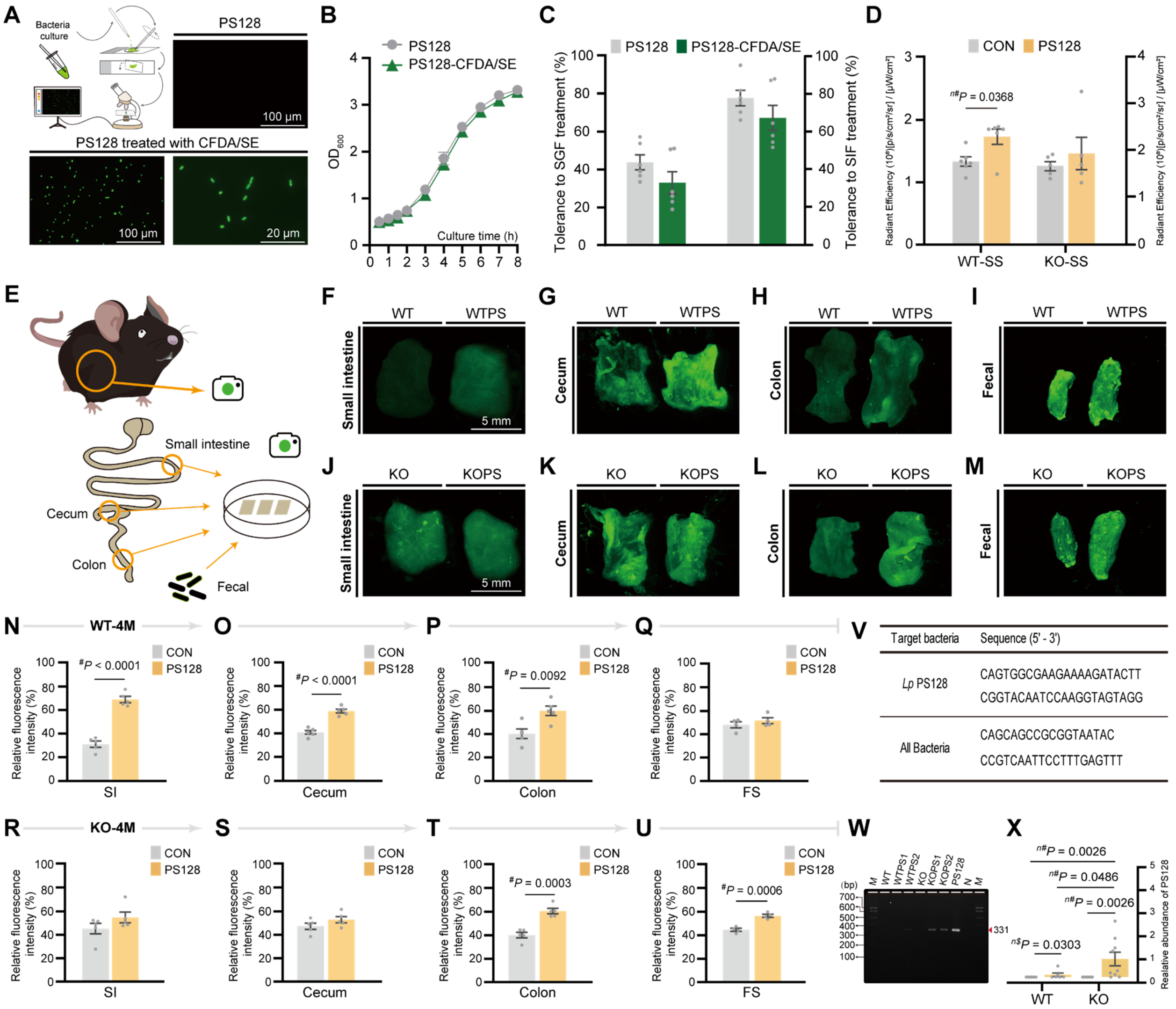


**Fig. S3 Localization and detection of the administered *Lactiplantibacillus* strain across mouse groups, related to Fig. 2. A** Schematic of CFDA/SE–labeled PS128 and representative fluorescence microscopy images. Scale bars, 100 μm and 20 μm. **B** Growth curves of CFDA/SE–labeled (green) and unlabeled (gray) PS128 at 37°C. **C** Tolerance of PS128 to simulated gastric fluid (SGF) and simulated intestinal fluid (SIF), expressed as percentages. **D** Biodistribution of PS128 in the stomach and small intestine (SS). Average radiant efficiency quantified from the SS, $n$ = 5–6. **E–M** Schematic of fluorescence imaging of intestinal tissues from adult *Fmr1* KO and wild-type (WT) mice with or without PS128 intervention (**E**). Small intestine (SI) (**F** and **J**), cecum (**G** and **K**), colon (**H** and **L**), and fecal surface (FS) (**I** and **M**), $n$ = 4–5. Scale bars, 5 mm. **N–U** Relative fluorescence intensity (%) of gut tissue fragments and feces. Small intestine (SI) (**N** and **R**), cecum (**O** and **S**), colon (**P** and **T**), and fecal surface (FS) (**Q** and **U**), $n$ = 4–5. **V–X** Quantification of PS128 in fecal samples. PS128-specific primers used for identification (**V**). PCR detection of PS128 in feces and pure culture. N, negative control; M, 100-bp ladder (**W**). Relative abundance of PS128 determined by RT-*q*PCR (**X**), $n$ = 6–9. Data are presented as mean ± S.E.M., and each dot represents an individual mouse. Statistical significance was determined by two-tailed Student's *t*-test ($^{\#}$), one-tailed ($^{n\$}$) or two-tailed ($^{n\#}$) Mann–Whitney U tests, as indicated. A $P$ value < 0.05 was considered statistically significant.

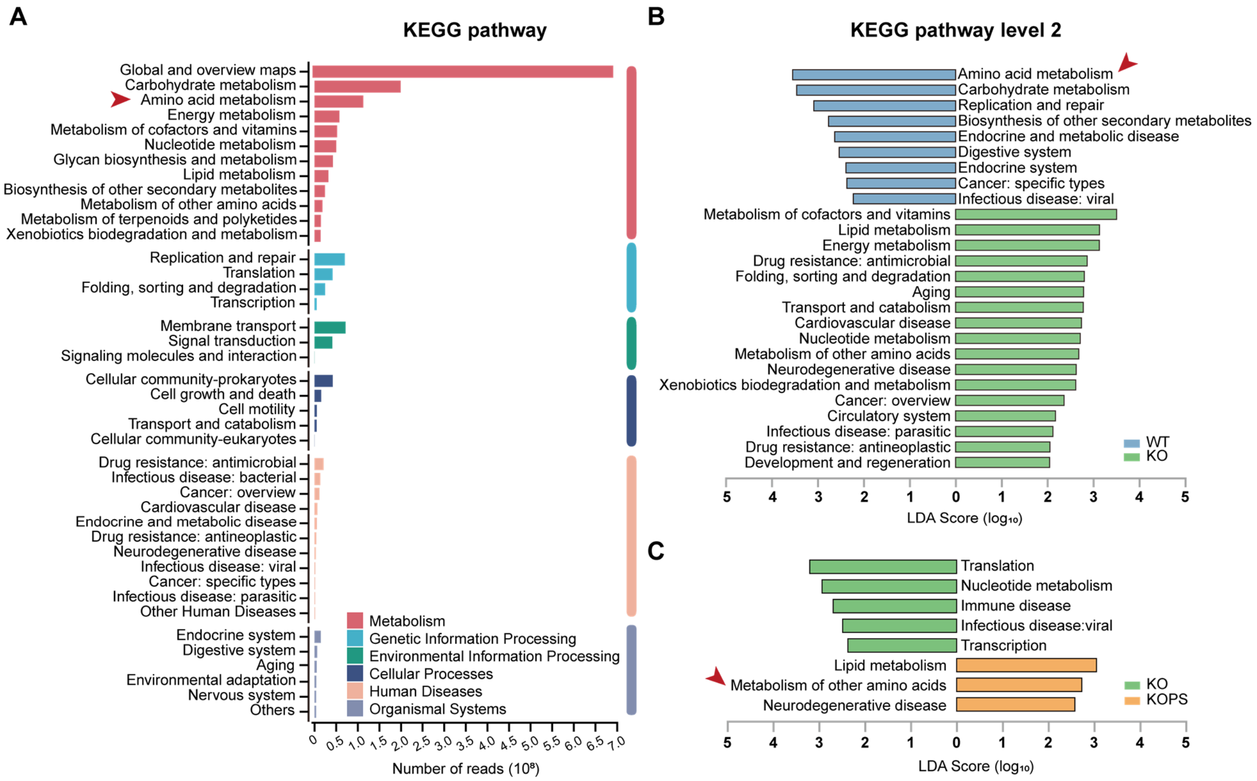


**Fig. S4 Enrichment analysis of KEGG pathways, related to Fig. 4.** **A** Enrichment map of Kyoto Encyclopedia of Genes and Genomes (KEGG) pathways. **B** and **C** Histograms of linear discriminant analysis (LDA) scores for differentially abundant KEGG level 2 pathways identified by linear discriminant analysis effect size (LEfSe) in comparisons between wild-type (WT) and *Fmr1* knockout (KO) mice (**B**) and between KO and PS128-treated KO (KOPS) mice (**C**). LDA score cutoff > 2.0, $P < 0.05$, $n = 11–13$ per mouse group. Red arrows denote pathways related to amino acid metabolism.

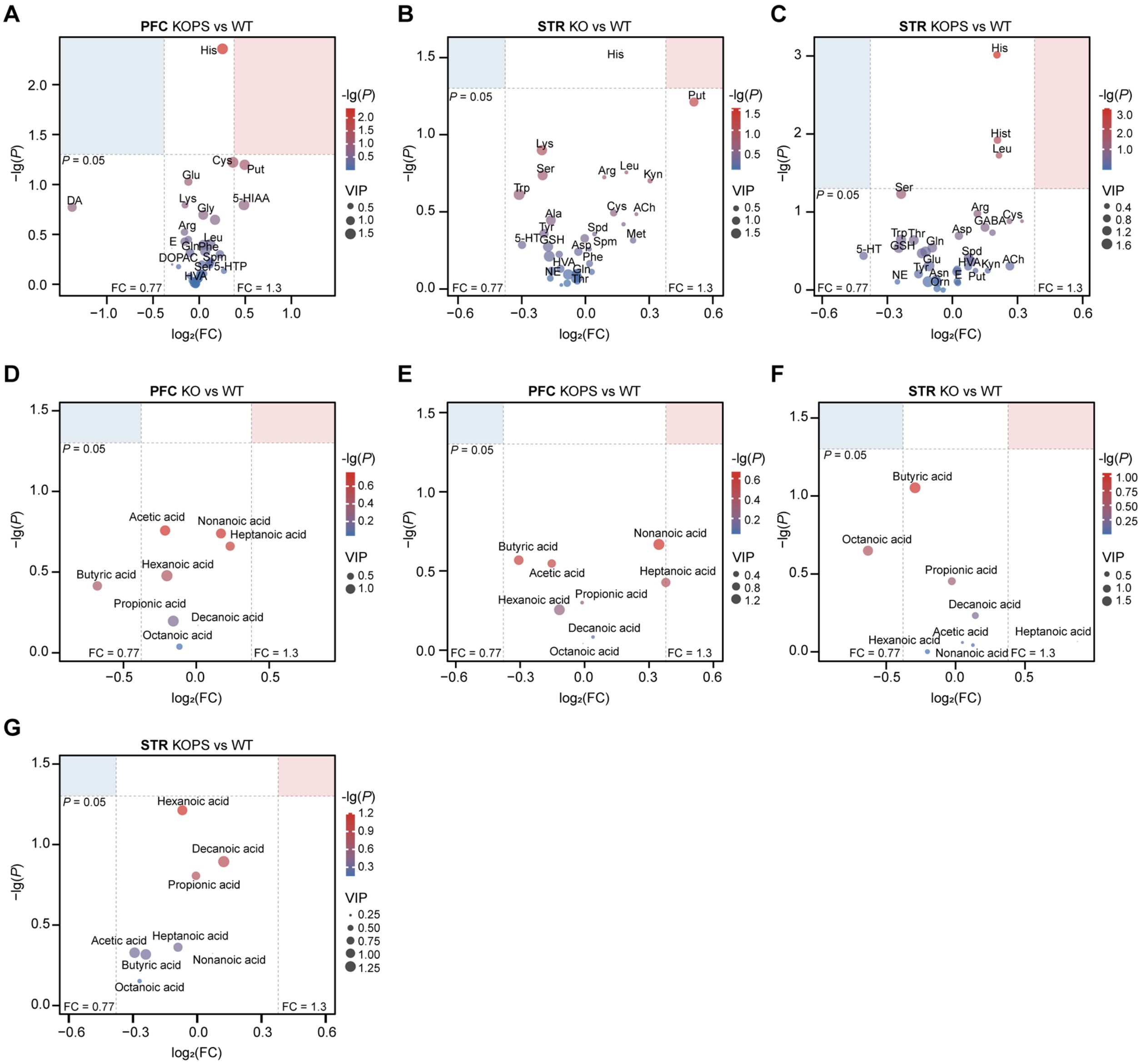


**Fig S5. Targeted metabolomic profiling of neuroactive metabolites and fatty acids in PFC and striatum, related to Fig. 5. A–G** Volcano plots showing differential neuroactive metabolites and fatty acids in the prefrontal cortex (PFC) and striatum (STR). Neuroactive metabolites in the PFC (**A**) and STR (**B** and **C**) comparing KOPS versus WT (**A**), KO versus WT (**B**), and KOPS versus WT (**C**). Fatty acids in the PFC (**D** and **E**) and STR (**F** and **G**) comparing KO versus WT (**D** and **F**) and KOPS versus WT (**E** and **G**). Colors indicate $-\log_{10}(P$ value), and bubble sizes represent variable importance in projection (VIP) scores. The x-axis denotes $\log_2$(fold change). FC, fold change. PFC, $n = 3$ biological replicates pooled from six individuals (18 mice per group). STR, $n = 4$ biological replicates pooled from three individuals (12 mice per group).

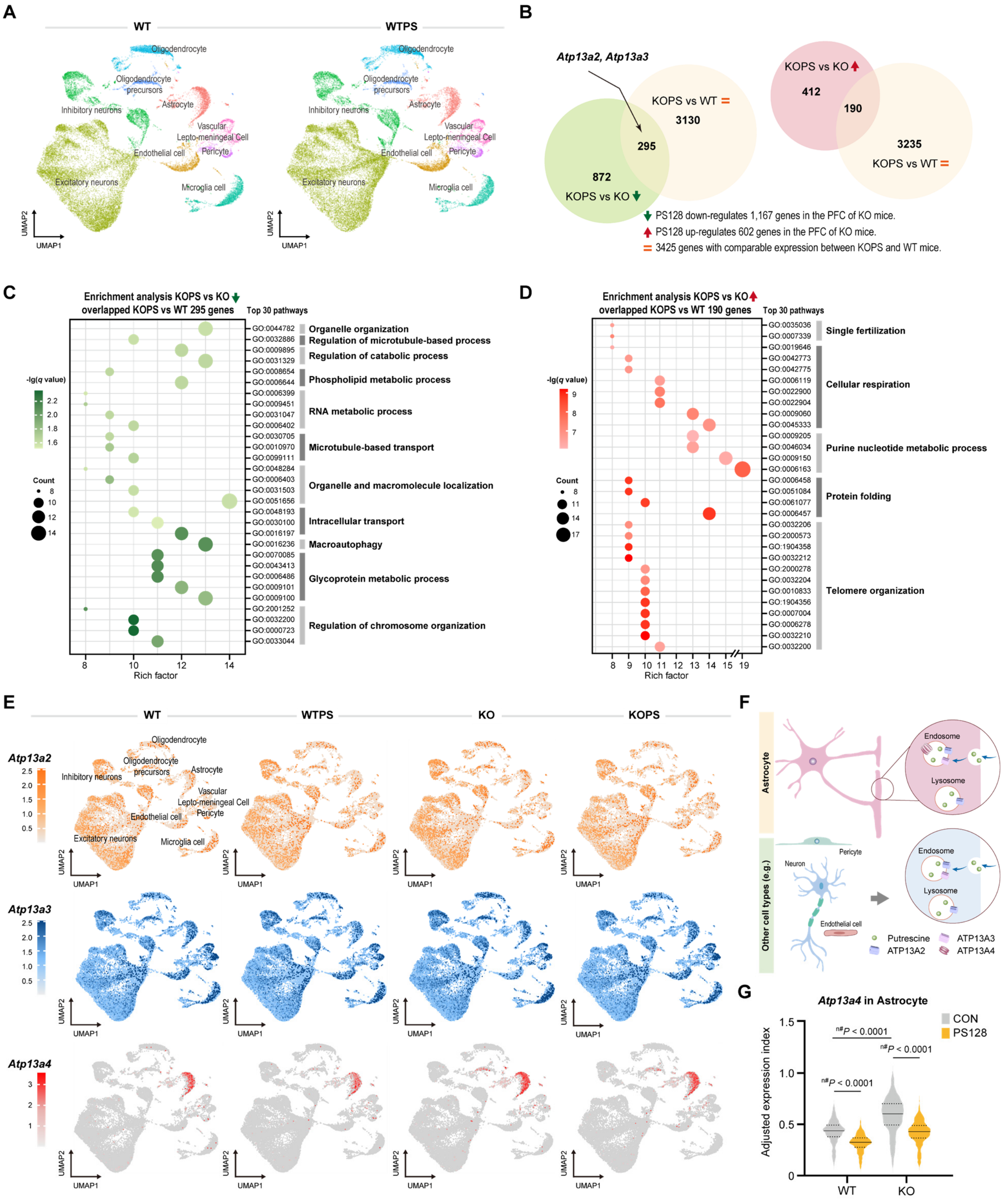


**Fig. S6 Transcriptomic changes in the prefrontal cortex following PS128 intervention in *Fmr1* KO mice, related to Fig. 5. A** UMAP projection of cell-type composition derived from single-nucleus RNA sequencing (snRNA-seq) of the prefrontal cortex (PFC) from 15-week-old mice. $n = 2$ biological replicates per group, each pooled from 3–4 individuals (7–8 mice per group). **B** Venn diagrams of differentially expressed genes (DEGs) identified by snRNA-seq analysis. DEGs were defined as $P < 0.05$ and expression in > 9% of PFC cells. A total of 1,167 genes were downregulated in KOPS relative to KO mice (green), 602 genes were upregulated (pink), and 3,425 genes showed similar expression between KOPS and wild-type (WT) mice (light yellow). Among these, 295 genes overlapped between the downregulated and WT-similar sets, and 190 genes overlapped between the upregulated and WT-similar sets. **C** and **D** Gene Ontology (GO) enrichment analyses of the 295 overlapping genes (**C**) and 190 overlapping genes (**D**) identified in (**B**). **E** UMAP feature plots showing normalized expression of *Atp13a2* (gray to deep orange), *Atp13a3* (gray to deep blue), and *Atp13a4* (gray to deep red) across WT, WTPS, KO, and KOPS mice. Cells with no detectable expression are shown in gray, and higher expression levels are indicated by increased color intensity. **F** Schematic illustration highlighting the cell-type distribution of *Atp13a2*, *Atp13a3*, and *Atp13a4* expression. **G** *Atp13a4* expression index in astrocytes. Astrocyte counts were 396, 261, 631, and 547 cells for WT, WTPS, KO, and KOPS groups, respectively. The *Atp13a4* expression index was calculated as the product of (i) the proportion of *Atp13a4*-positive astrocytes (cells with detectable expression) and (ii) the mean normalized *Atp13a4* expression level among those positive cells, based on Seurat-processed snRNA-seq data. Statistical significance was determined using a two-tailed Mann–Whitney U test ($^{n\#}$). A $P$ value < 0.05 was considered statistically significant.

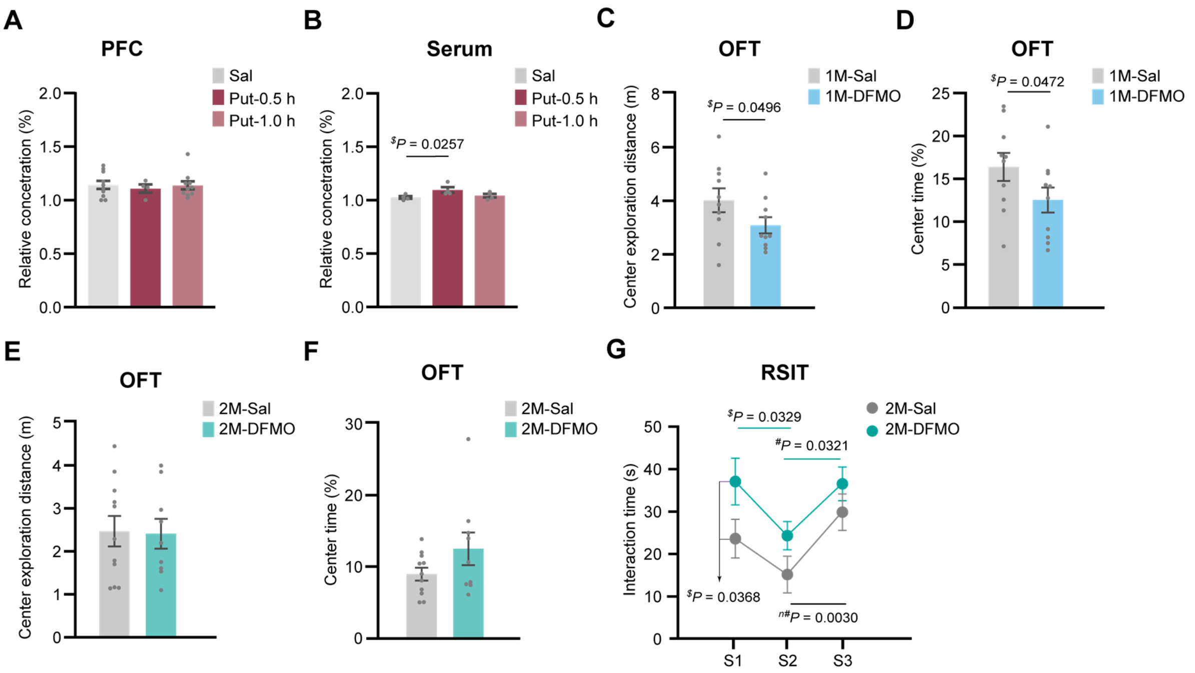


**Fig. S7 Putrescine imbalance and autism-like behaviors in mice, related to Fig. 6. A** and **B** Relative putrescine levels in the prefrontal cortex (**A**) and serum (**B**) of wild-type (WT) mice, $n$ = 4–10. **C–F** Center exploration distance (**C** and **E**) and time spent in the center area (**D** and **F**) in the open-field test (OFT), $n$ = 9–11. **G** Interaction time with familiar or novel mice across three stages of the reciprocal social interaction test (RSIT). S, stage. Data are presented as mean ± SEM, and gray dots indicate individual mice. Statistical significance was determined using one-tailed ($^{\$}$) or two-tailed ($^{\#}$) Student's $t$-tests, or two-tailed ($^{n\#}$) Mann–Whitney U test, as appropriate. A $P$ value < 0.05 was considered statistically significant.
